# Machine-Learning-Based Olfactometry: An Auxiliary System for Human Assessors in Olfactory Measurement

**DOI:** 10.1101/2022.04.20.488973

**Authors:** Liang Shang, Chuanjun Liu, Fengzhen Tang, Bin Chen, Lianqing Liu, Kenshi Hayashi

## Abstract

Although gas chromatography/olfactometry (GC/O) has been employed as a powerful analytical tool in odor measurement, its application is limited by the variability, subjectivity, and high cost of the trained panelists who are used as detectors in the system. The advancements in data-driven science have made it possible to predict structure-odor-relationship (SOR) and thus to develop machine-learning-based olfactometry (ML-GCO) in which the human panelists may be replaced by machine learning models to obtain the sensory information of GC-separated chemical compounds. However, one challenge remained in ML-GCO is that there are too many odor descriptors (ODs) being used to describe the sensory characteristics of odorants. It is impractical to build a corresponding model for each OD. To solve this issue, we propose a SOR prediction approach based on odor descriptor clustering. 256 representative ODs are firstly classified into 20 categories using a co-occurrence Bayesian embedding model. The categorization effect is explained according to the semantic relationships using a pre-trained Word2Vec model. Various molecular structure features including molecularly parameters, molecular fingerprints, and molecular 2D graphic features extracted by convolutional neural networks, are employed to predict the aforementioned odor categories. High prediction accuracies (Area under ROC curve was 0.800±0.004) demonstrate the rationality of the proposed clustering scenario and molecular feature extraction. This study makes the ML-GCO models much closer to the practical application since they can be expected as either an auxiliary system or complete replacement of human panelists to perform the olfactory evaluation.

## 1. Introduction

Gas chromatography/olfactometry (GC/O) is a key technique that integrates the separation of volatile compounds using a gas chromatograph (GC) with the detection of odor by employing human assessors as olfactometers [1-4]. Contributed by mass spectrometry (MS) analysis and human assessors, GC/O can provide not only the molecular information of complex odor mixtures, but also sensory information of specific odor-active components. Therefore, it has been applied as a critical analytical instrument in various filed, such as food, cosmetics, agriculture, and the environment [5-8].

Although human assessors play a very important role, the promotion and application of GC/O measurement in the meantime are also limited by the limitations of human evaluation. The GC/O measurement generally requires multiple people to detect and evaluate volatile compounds by sniffing GC effluent components in real-time [9]. Therefore, the subjectivity of panelists at the intra- and inter-individual levels is inevitable. The high cost of training and employing human panelists is also a great problem. In addition, limited by the resolution of the GC instrument as well as the olfactory threshold of human olfaction, the odor-active compounds may be ignored during the GC/O measurement. Recently, many auxiliary software modules, such as AroChemBase (Alpha MOS Co. Ltd.) and GC/MS Off-Flavor Analyzer (Shimadzu Co. Ltd.), have been developed as supplements to GC/O analysis. These auxiliary systems work well in improving the accuracy of the evaluation as well as in reducing the manual labor of human assessors.

It is well known that the number of odorants with known descriptors is limited despite having long years of accumulation in the field of flavors and fragrances. The number of odorants with odor descriptors (around 2000) that are registered in those auxiliary systems is far less than that of the total odorant pool (around 44,000) [10]. These existing systems may not meet the requirements of GC/O measurement since most of the odor-active compounds still remain unknown to us. This problem may persist for a long time given the difficulty of olfactory measurement. Recently, the use of artificial intelligence (AI) to support analytical purposes has been a potential technology for overcoming the drawbacks of traditional analytical methods [11, 12].

The precise relationship between molecular structure and odor perception, termed the structure-odor relationship (SOR), has attracted considerable attention from the aspect of data science [13, 14]. Great efforts have been made in ODs identification based on various molecular features, such as physicochemical parameters, bio-inspired olfactory model, and MS [15]. Moreover, various deep neural network models (DNNs), machine learning (ML) classification frameworks, and topic models have been developed to express the SOR [16-18]. More recently, Debnath and co-workers proposed a model to predict perceptual scent impressions from MS of odorants under an imbalanced dataset [19, 20]. In addition, Mayhew et. al. developed an odorless identification model based on transport features with an extremely high accuracy [21]. The abovementioned studies indicated that the SOR would be solved by ML algorithms and chemometrics. Therefore, our research team previously developed a method for conducting ML-based GC/O (ML-GCO), in which olfactometry detection can be done by an ML classifier, thus reducing the need for a human panelist [22].

It should be mentioned that, however, there are many problems needed to be overcome before the practical application of ML-GCO. For example, the dimension of odor space remains unknown and it is not yet clear on the primary dimensions of olfactory like vision or gustatory [23]. Odors are highly complex and people are known to disagree regarding their linguistic descriptions of smell sensations. The number of odor compounds is estimated at over 400,000 which are described by several hundred odor descriptors [24]. In most reported work, only a small number of typical, common odor descriptors have been included, leaving the majority of them undefined [22, 25, 26].

Many researchers have demonstrated that odor descriptors are inherently connected in the odor structure space. For example, carbon-chain length and functional groups have been confirmed as critical factors in odor perception [27-29]. This means the odor descriptors can be clustered according to their similarity evaluations and thus cover the odor descriptor space as many as possible. One challenge in the prediction of ODs is defining the primary dimensions of the olfactory perceptual space [30]. In addition, odors are highly complex, and people are known to disagree regarding their linguistic descriptions of smell sensations [31-33]. Although most ODs can be predicted with high precision, infrequent smells, such as apricot and chocolate, were shown to difficult to be predicted [25]. Yet, previous studies have demonstrated that odor descriptors are inherently connected in perceptual space [26]. It indicated that co-occurrence of ODs from odor database would be a reasonable direction to find odor categories. Recently, Pandey and co-workers proposed a vibration-based biomimetic odor classification method based on the vibrational spectrum of an odorant molecule [34]. However, only 20 odorant molecules were considered for analyzing odor perceptual classes, which would not enough to offer insight into the olfaction space. Therefore, this issue could be addressed via a rational relationship for OD similarity evaluations and clustering using a large odor database.

In response to the above problems, ODs embedding and clustering approaches are proposed. A schematic of the data-processing method is illustrated in Fig. 1. ODs from different data sources are collected and merged, and Bayesian co-occurrence embedding (BCE) was employed for odorants and ODs embedding. Based on the distance between odorant vectors and OD vectors, ODs can be calibrated and relabeled. And then hierarchical clustering analysis (HCA) was performed on the embedded OD vectors to investigate the internal relationship between ODs and their clustering result is discussed. We show that a total of 256 ODs were clustered in 20 categories, and most of the categories can be supported by previous research. In addition, the structure parameters of odorant molecules are employed for training ML models to verify the rationality of the proposed clustering scenario. Results show that the smell categories can be predicted by the models established by molecularly structure features of odorants successfully (Area under ROC curve: 0.800±0.004, precision: 0.595±0.004, recall: 0.721±0.003 and F-score: 0.570±0.007, p<0.0001 Wilcoxon test). It indicated that the smell clusters proposed in this study are supported by the SOR. The proposed odor category is not only expected as a novel research direction for developing ML-GCO, but also applies a reference for understanding the biology of olfaction. This study makes the ML-GCO models much closer to the practical application since they can be expected as either an auxiliary system or complete replacement of human panelists to perform the olfactory evaluation.

**Fig. 1.**
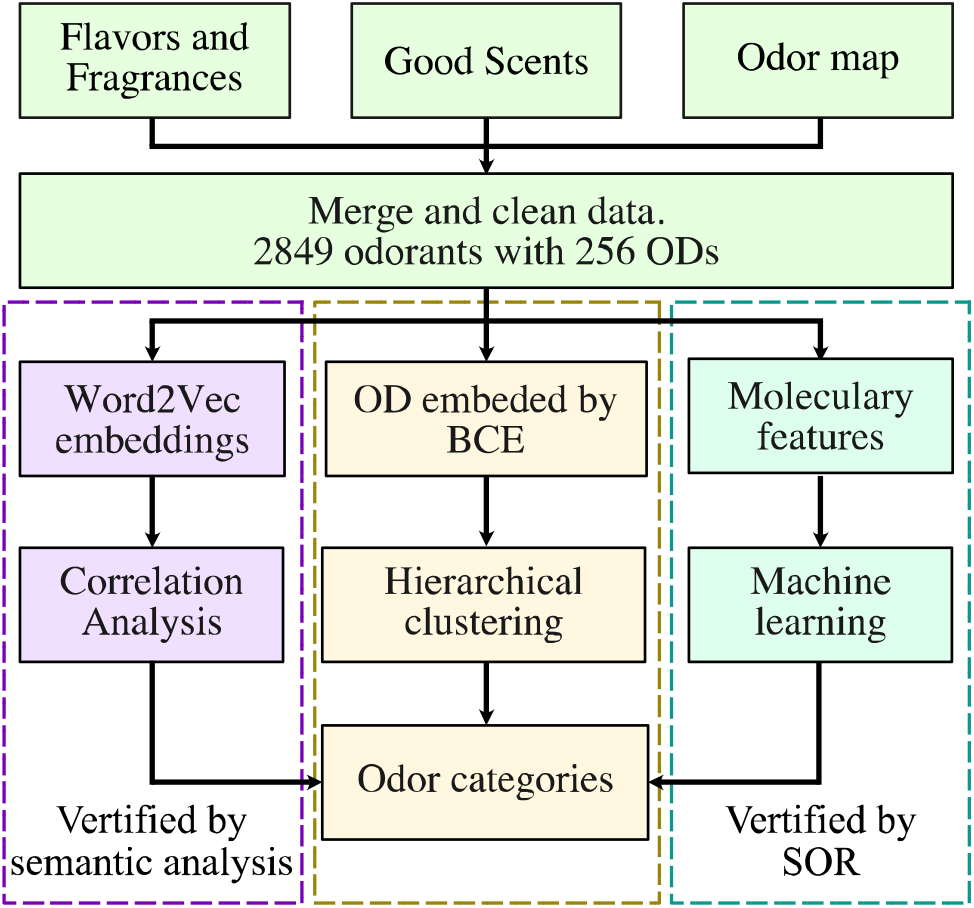
Data processing diagram of odor descriptor clustering analysis.

## 2. Material and methods

### 2.1 Data preparation

To understand the internal relationships between smell percepts, we collected the chemical abstracts service (CAS) data of odorants and their odor perceptions via both web scraping and manual methods. We used three publically available odor databases, including the Flavors and Fragrances database (Sigma-Aldrich) [35], the Good Scents database [36], and the Odor Map database [37]. Detailed information on odor databases used in the present study is summarized in Table 1. Based on CAS number, the simplified molecular-input line-entry system (SMILES) data were collected from PubChem (https://pubchem.ncbi.nlm.nih.gov) [38]. Using RDKit software (http://www.rdkit.org) [39], molecular structure images, molecular parameters, and molecular fingerprints were obtained for further analysis. In the present study, chemicals without odor descriptors or those considered “odorless” were not considered. Finally, 2849 molecules with 510 ODs were collected and analyzed. All of the ODs in the database are listed in Table S1.

**Table 1.** Summary of odorant databases used in present study.

| Data base | # odorants | Description |
| --- | --- | --- |
| Flavors and Fragrances | 1026 | An odor database of flavor and fragrance proposed by Sigma Aldrich. |
| Good Scents | 3673 | An odor database of flavor, fragrance, food and cosmetic industries. |
| OdorMapDB | 321 | An odor database of odorants and their olfactory bulb responses (odor maps). |

### 2.2 Bayesian co-occurrence embedding

To normalize the labels for odorants from all databases, BCE was introduced for OD vector embedding [40]. A brief description of BCE is illustrated in Fig. 2. Based on BCE, a preference matrix for each odorant can be generated. A plus sign (+) indicates comparisons with OD_d_, OD_j_ is the label for the odorant_i_, a minus sign (-) indicates the opposite, and a question mark (?) indicates the unknown value that is estimated by the BCE algorithm. Therefore, the individual probability that an odorant prefers OD_d_ to OD_j_ was defined as:

**Fig. 2.**
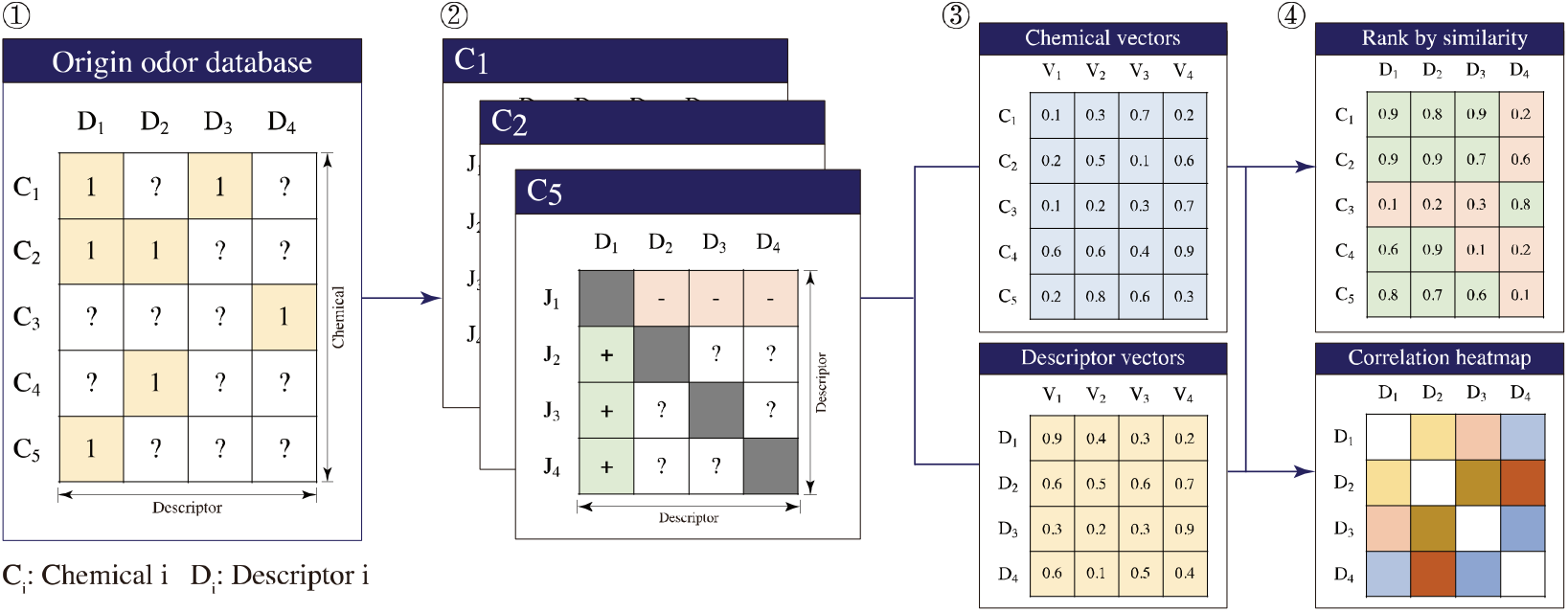
Odor descriptor embedding and calibration using the Bayesian co-occurrence embedding (BCE) method.

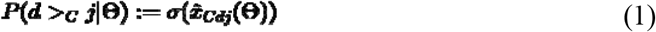

where ***σ*** is the logistic sigmoid function, **θ** is the parameters of the BCE model, and 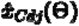 is the score indicating the degree to which chemical C prefers OD_d_ rather than OD_j_. According to maximum posterior estimation, the generic optimization criterion for each odorant could be estimated as follows:

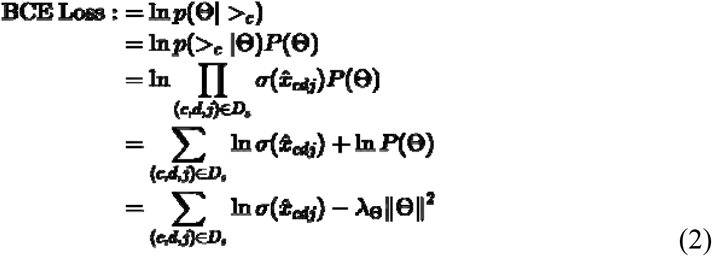

where **λ**_**θ**_ are model-specific regularization parameters. Therefore, the gradient of the loss function with respect to the model parameters is:

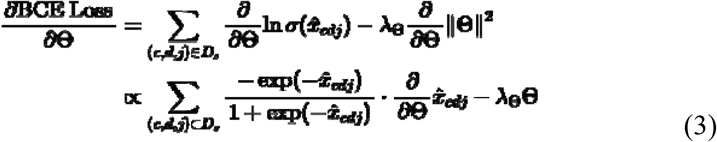

Finally, the model parameters (**θ**) could be updated using the assigned learning rate ***η*** and stochastic gradient descent as follows.

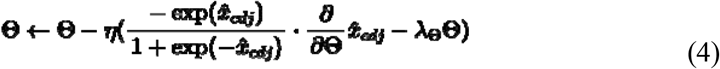

Unlike k-nearest neighbor (kNN) collaborative filtering or matrix factorization (MF), BCE applies a Bayesian optimization criterion to generate odorant similarity rankings based on pairs of ODs (i.e. the odorant-specific order of two ODs). As an offline embedding method, Bayesian optimization has advantages over the standard learning techniques for MF and kNN [41, 42]. In the present study, the relabeling results produced by BCE were evaluated via the normalized discounted cumulative gain (NDCG).

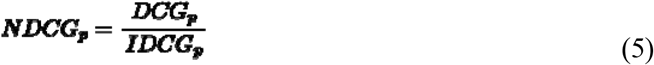

where the discounted cumulative gain (DCG) and ideal discounted cumulative gain (IDCG) were calculated as:

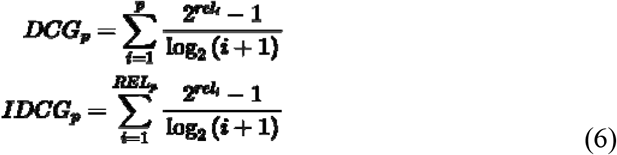

### 2.3 Molecular graphic feature extraction

To extract the necessary numerical features from the molecular structure images, we considered four types of pretrained convolutional neural network (CNN) frameworks, including VGG-16, Restnet, Densnet, and Alexnet. Detailed information for these CNNs is given in these papers [43-46]. In the present study, the CNNs were only applied as molecular-graphic feature extractors for the SOR model calibration.

### 2.4 SOR model calibration

The dataset for the odor category prediction was a typical imbalanced data set because the class distribution of the positive samples (minor samples with specified odor categories) and negative samples (major samples with non-specified odor categories) was not uniform. Inspired by Mordelet’s research, we considered bagging classifiers to be a feasible method for learning with an imbalanced data set [47]. Detail description of transductive bagging learning is presented in the support information. In the present study, the sample pool was divided into training and test sets with a 3:1 ratio via random stratified sampling. In addition, the number of bootstraps was set as 100 and the subsample number was the same as that for the positive numbers. Considering the sample size, we employed GBDT and GLVQ to predict the odor clusters in the present study [48, 49]. More introduction to these models is described in the support information. Finally, the optimal feature extractor and model combination was determined by considering the area under the ROC curve (AUC-ROC), precision, recall, and F1-score of the test set, respectively. Detailed information for these metrics is presented in the support information materials. In the present study, data and models were processed and analyzed using Python (ver. 3.9.0) and R (ver. 4.1.1).

### 2.5 Word2Vec embedding model

To investigate the semantic internal relationships between ODs for each odor category, we used Google’s pre-trained Word2Vec model to create semantic presentations for ODs. The model, which contains 300-dimensional vectors for 3 million words and phrases, was trained using the Google News dataset. Two types of model architectures, including the continuous bag-of-words (CBOW) model and the continuous skip-gram model, were developed for learning latent presentations for words. More detailed information can be found here [50]. The proposed model has been confirmed to perform better than previous techniques based on different types of neural networks. In addition, the proposed vectors provide the state-of-the-art performance for measuring semantic word similarities.

## 3. Results and Discussion

### 3.1 Odor perception embedding and calibration

In the present study, the idea of co-occurrence in BCE was introduced for odor perception calibration. Using the BCE method, ODs can be embedded as numerical vectors. According to the cosine similarity between the odorant and OD vectors, the top 20 nearest ODs were considered as candidates for each odorant. As a critical factor, optimization of the embedded dimension is an important consideration. The normalized discounted cumulative gain for the top 20 ODs (NDCG@20) under different embedded dimensions is shown in Fig. S1. The NDCG@20 increased with the number of embedded dimensions. Embedded vectors with 64 dimensions performed significantly better than embedded vectors with 32, 16, and 8 dimensions (p<0.001), and were not significantly different from embedded vectors with 128, 256, 512 dimensions. Considering accuracy and computational efficiency, we selected 64 as the optimal number of embedded dimensions for the BCE in the present study. The distribution of ODs before and after calibration is shown in Fig. S2, and detailed information is given in Table S1. In summary, the sample size of the ODs increased after calibration in 70.58 % of cases. In this type of statistics, the sample size should always be more than 20. Using the BCE algorithm, the sample size increased to more than 20 in 61 ODs (11.96 %), which were then considered for further analysis. Finally, 265 ODs (51.96 %) were selected for afterward clustering analysis in the present study.

### 3.2 Clustering characterization for odor descriptors

To quantify the inner relationships between ODs, we performed hierarchical clustering based on the Euclidean distances between the embedded vectors of smell descriptors. Because the basis of olfaction has not yet been established, we turned to previous research to explain our clustering results. The well-known Dravnieks [51] and DREAM [52] datasets include 19 types of descriptors: the scent of a bakery, sweet, fruit, fish, garlic, spices, cold, sour, burnt, acid, warm, musky, sweaty, ammonia/urinous, decayed, wood, grass, floral, and chemical [53]. Based on an analysis of previous research [54, 55], we considered 20 to be a reasonable number of clusters. The results were organized and depicted using dendrograms, as shown in Fig. 3 and Fig. S3, S4. In the clustering results, the descriptors with semantic similarity were almost always grouped in the same class. Specifically, cluster-1 was composed of sweaty-like and fish-like descriptors, which were considered unpleasant odors. ODs related to the scent of a bakery were clustered in cluster-2, and burnt-like descriptors were present in cluster-4 and cluster-12. Most groups, including cluster-5 (milky-like), cluster-6 (spicy-like), cluster-7 (musky or green-like), cluster-10 (wine or ester-like), cluster-11 (fruit acid-like), cluster-13 (cold or fresh-like), cluster-14 (garlic-like), cluster-15 (fruit-like), cluster-16 (floral), cluster-17 (cold-like), cluster-18 (musky and wood-like), and cluster-20 (wood-like), were supported by the core smell descriptors proposed in previous research [56].

**Fig. 3.**
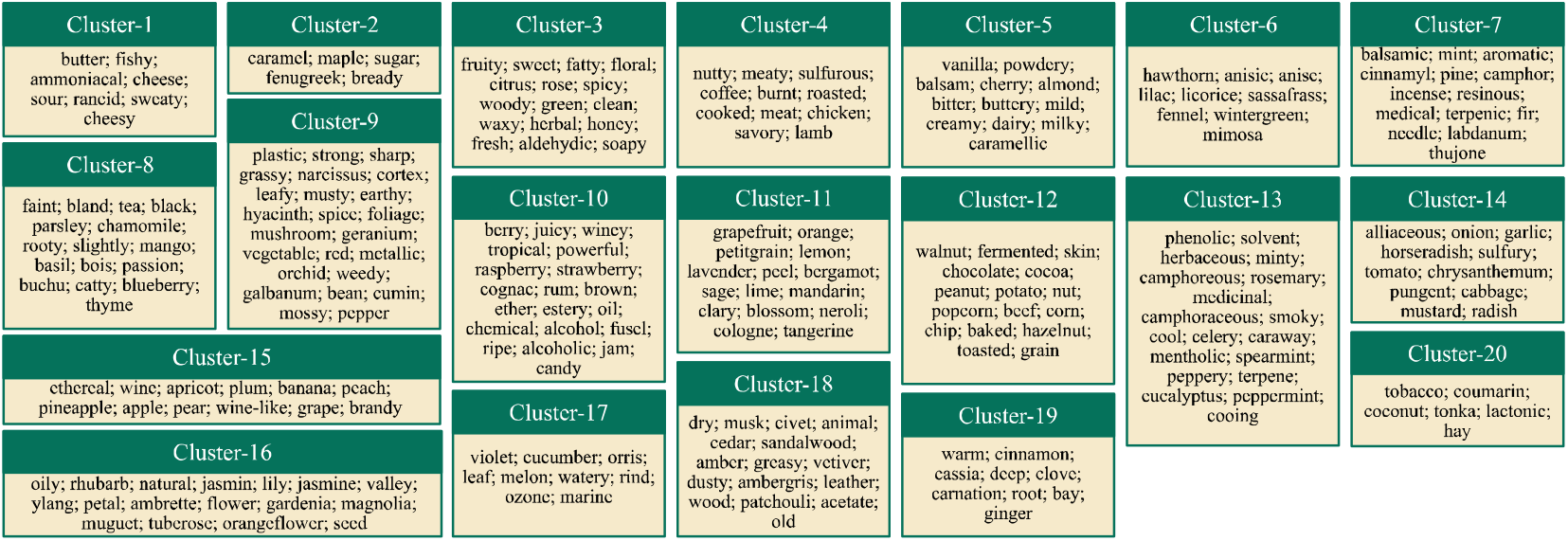
A summary of odor descriptor clustering based on the similarities between BCE embedded vectors.

Although most of the descriptors with similar linguistic meanings were clustered in the same group, special cases were also observed in some clusters. For cluster-3, the ODs included fruity, sweet, floral, etc., and so we regarded this as the sweet or fruit-like group. However, some descriptors related to plants, such as spicy, woody, herbal, and green, were also present in cluster-3. In addition, we found tomato, chrysanthemum, rotten cabbage, and radish in cluster-14, and labeled this as the garlic-like group. To investigate the reasons for this “mis-clustering” of smell descriptors, the odorants contained in these ODs were extracted and analyzed. For example, the odor of methyl mercaptan (CAS: 74-93-1) is reminiscent of garlic or rotten cabbage, while Erucin (CAS: 4430-36-8) and Berteroin (CAS: 4430-42-6) are labeled as the odors of cabbage and radish. The link between “garlic” and “rotten cabbage”, and between “rotten cabbage” and “radish” indicated that these three descriptors could be clustered together, which can explain the presence of cabbage and radish in garlic-like clusters. In addition, Methyl propyl disulfide (CAS: 2179-60-4) is labeled as “radish”, “mustard”, “tomato”, “garlic”, etc., and Ethyl methyl sulfide (CAS: 2179-60-4) is labeled as “garlic”, “tomato”, “rotten cabbage”, etc. Thus, the odor semantic database showed an internal connection between the smell of tomato and that of garlic, which could be explained by the latent relationships between the ODs. The above-mentioned analyses demonstrated that semantic descriptor clustering based on embedded vectors using the BCE method is a reasonable approach for understanding internal affiliations. Even though most of the clusters could be explained by common sense or previous research, some clusters, such as cluster-8, were not found to belong to any ‘well-known’ smell categories proposed by previous studies. In cluster-8, descriptors such as tea, root, basil, thyme, and buchu could be regarded as woody or herbal. Furthermore, fruit-like (mango, blueberry, passion fruit) and mild descriptors (faint, bland, slightly) were also clustered here. The plants associated with these ODs, such as tea or basil, are mild and herbaceous, which could explain this clustering result. In cluster-10, most of the descriptors were related to alcohol, such as wine-like, powerful, rum-like, alcoholic, etc. Wine flavor descriptors, such as berry, juicy, jam, candy, ether, and ester, were also identified. Consequently, cluster-10 was considered to have the aroma of wine. Chemical-like descriptors, including oil, chemical, and fusel, were also clustered here, which indicated that cluster-10 comprised multiple types of smell perceptions. Interestingly, unpleasant smell descriptors, such as sweaty/fish-like (cluster-1) and garlic-like (cluster-14), could be easily clustered, which confirmed that flavor-like impressions were regarded as more complex than unpleasant perceptions.

### 3.3 Mapping odor descriptors in t-SNE space

To investigate the internal relationships between odor categories, we visualized the data using Barnes-Hut t-distributed stochastic neighbor embedding (t-SNE) as an unsupervised low dimension presentation method. Detail explanation of the t-SNE method is described in support information. As illustrated in Fig. 4, odor descriptors from 20 categories were mapped in t-SNE space via manifold embedding (an interactive version is at https://shangliang0225.shinyapps.io/molodorperception.). The details for the embedded data calculated using t-SNE are presented in Table S2. The analysis illustrated that ODs from cluster-1 (sweaty or fish-like), cluster-2 (bakery-like), cluster-3 (woody or herb-like), cluster-7 (musky or green-like), cluster-11 (fruit or acid-like), cluster-12 (burnt-like), and cluster-14 (garlic-like) were clustered together. Additionally, descriptors from cluster-4 (burnt-like), cluster-13 (cold or fresh-like), cluster-16 (floral), and cluster-20 (wood-like) were clustered in multiple groups. For cluster-4, we found that part-1 (including chicken, lamb, and savory) was on the left of the t-SNE map, and part-2 (including nutty, meaty, sulfurous, coffee, burnt, roasted, cooked, and meat) was on the right of part-1. In addition, we found that beef was located near part-1 of cluster-4, which could be explained by the meat-like smell of beef. Smoky was located near part-2 of cluster-4, which was close to perceptions of burnt or roasted. In summary, plants or herb-related smell perception categories, such as cluster-6, cluster-7, cluster-8, cluster-13, and cluster-19, were located at the top area of the t-SNE space. In addition, fruity or alcohol-like categories (including cluster-10, cluster-11, and cluster-15) were neighbors, and were below the plant of herb-related categories. This demonstrates that the relationships between categories can be illustrated in t-SNE space, and that their locations can also be explained by their semantic meanings.

**Fig. 4.**
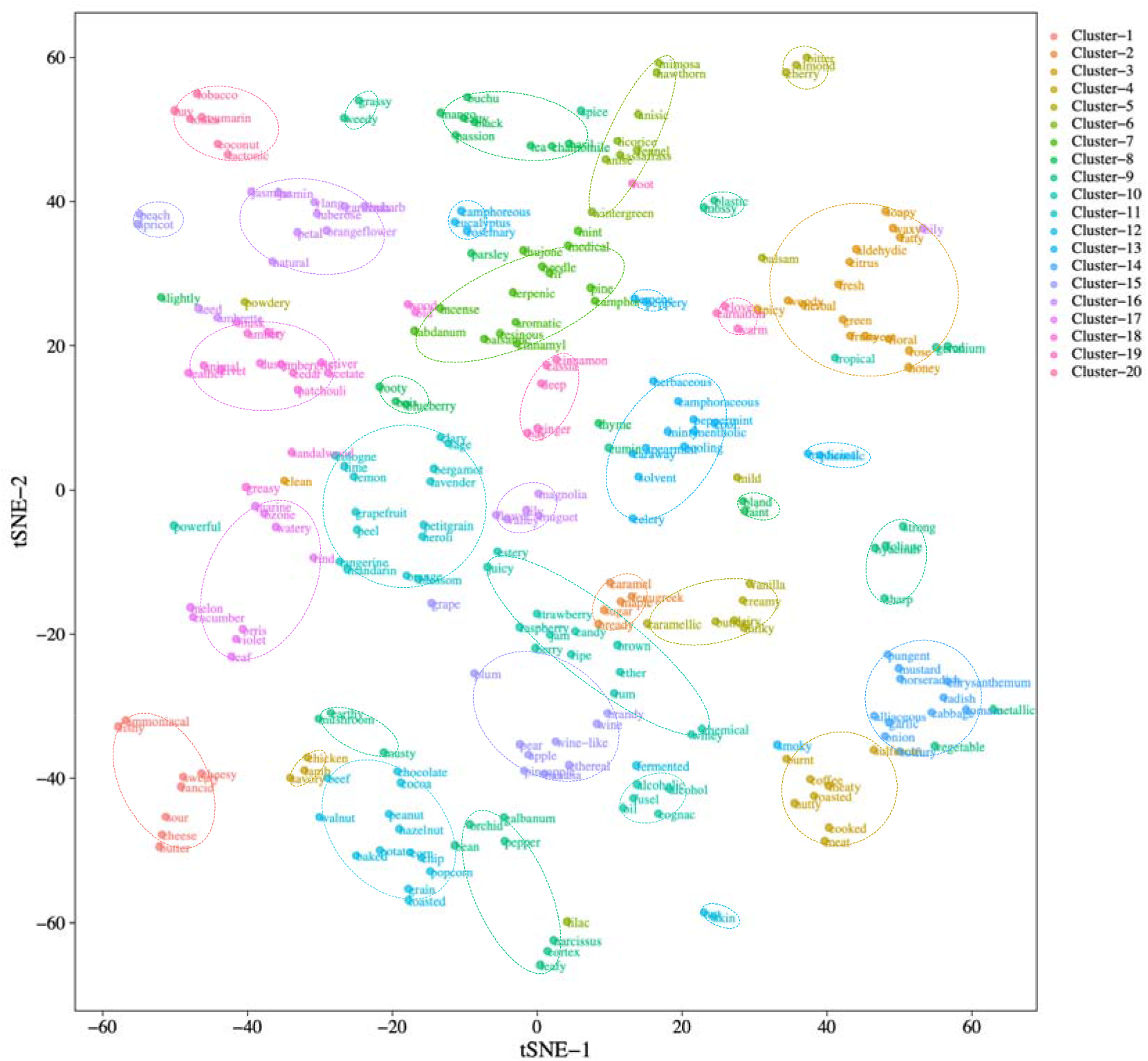
Odor descriptor clustering generated in the SOR space using the t-SNE method. An interactive version is available at https://shangliang0225.shinyapps.io/molodorperception.

### 3.4 Comparison with Word2Vec semantic embeddings

Odor descriptors are not only used to express odor feelings, but are also applied in daily written communication. To explore the pure semantic relationships for each category, we conducted a linguistic analysis. We employed a pre-trained Word2Vec model provided by Google as a feature extractor to generate word vectors containing 3 million words based on roughly 100 billion words from a Google News dataset. Correlation box plots for odor categories are given in Fig. 5 and Table S3. Correlation heat maps and their distributions are presented in Fig. S5 and Fig. S6. According to the analysis, cluster-14 (garlic-like, 0.479±0.128, p<0.0001) and cluster-6 (spicy-like, 0.443±0.0674, p<0.001) had higher internal correlations than the other odor categories, which indicated that these unpleasant perceptions had closer internal relationships in Word2Vec space. In contrast, some odor categories, such as cluster-10 (0.187±0.136, p<0.0001), cluster-8 (0.213±0.165, p<0.01), cluster-9 (0.217±0.139, p<0.05), and cluster-19 (0.227±0.174, p<0.0001), had lower correlations than the others. As mentioned above, cluster-10, cluster-8, and cluster-19 could not be defined as any ‘well-known’ smell categories. Furthermore, cluster-9 was composed of multi-odor perception categories, such as herb-like and chemical-like. However, the correlations between most odor categories were lower than 0.6, which demonstrates that the co-occurrence of terms in the text was not exactly the same as that of odor semantic descriptors.

**Fig. 5.**
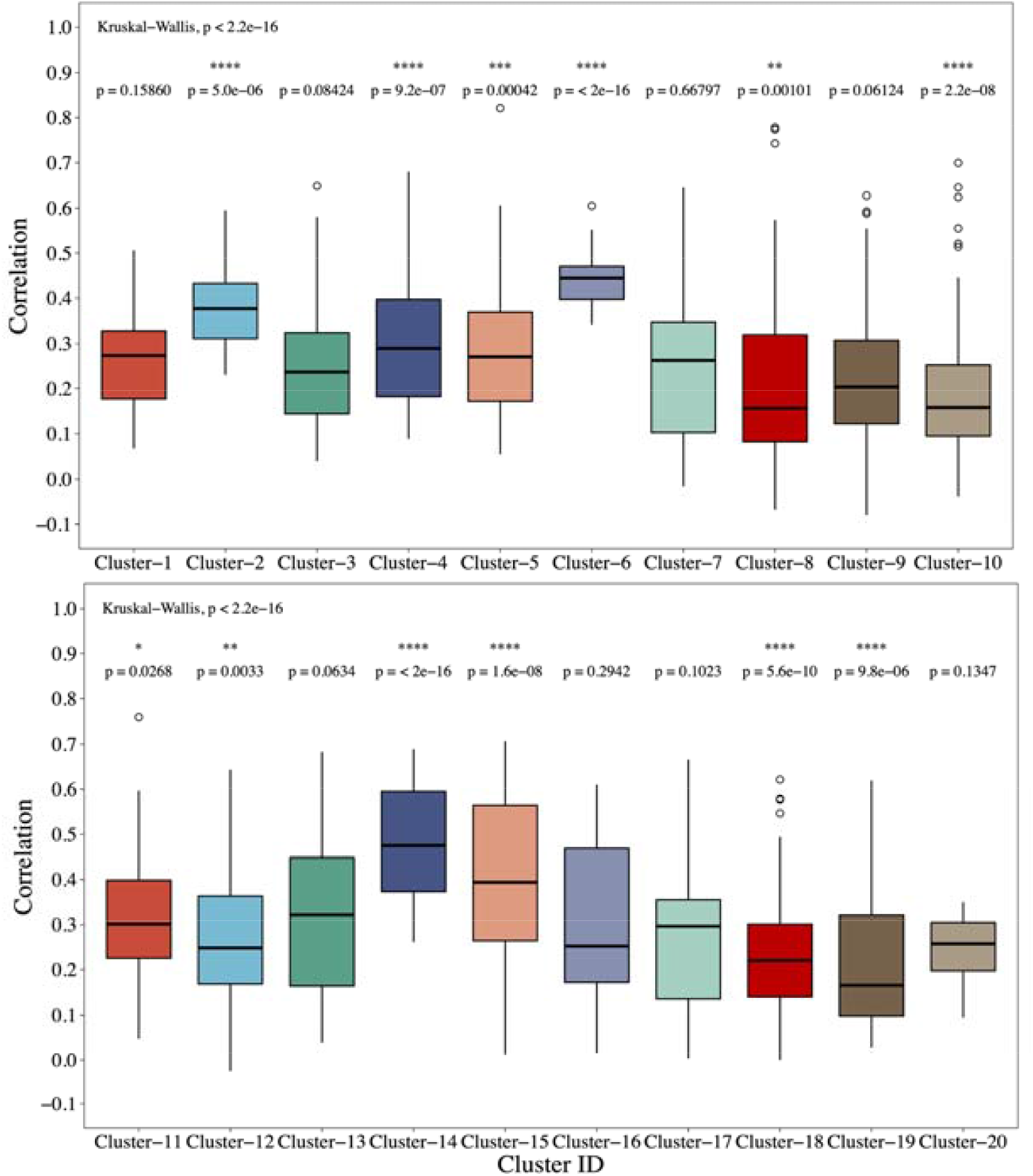
Correlation box plots of odor descriptor Word2Vec embeddings from odor perception categories. Results were evaluated using the nonparametric Wilcoxon signed-rank test.

### 3.5 Distribution of odor perception category labels

Before calibrating the odor category identification model, we first investigated the distribution of the samples. The sample distribution for each odor category is shown in Fig. S7a, which indicates that the sample sizes for the odor descriptors were clearly distinct and imbalanced. Fig. S7b illustrates the statistical distribution for the number of clusters of odorants. It demonstrates that most of the odorants belonged to more than two clusters, which can be explained by the complexity and ambiguity of odor perception.

### 3.6 Odor perception category identification models

To verify the rationality of the proposed clustering scenario, we used the molecular structure features to predict the aforementioned odor categories. To assess the potential of molecular feature extraction via four types of pre-trained CNN models, we used molecular parameters (MPs) and molecular fingerprints (FPs) to identify the clusters for odorants, applied GLVQ, and GBDT classification methods, and then compared and evaluated the results.

To calibrate the GLVQ models, the maximum number of training iterations, and the number of prototypes per class were set to 5000 and 10, respectively. The overall AUC, precision, recall, and F-score of the GLVQ models under the features extracted by CNNs, molecular parameters, and molecular fingerprint data sets are shown in Fig. 6, and the detailed predictions of the accuracies for each odor cluster are presented in Fig. S8 and Table 2. For cluster-2, cluster-13, and cluster-15, the VGG produced better results than the other feature extraction methods. However, the features extracted by Restnet did a better job in identifying most clusters. In summary, the Restnet model produced a significantly better average AUC (0.742±0.006), precision (0.580±0.003), recall (0.691±0.005), and F-score (0.548±0.009) than the other models (p<0.001).

**Table 2.** Odor cluster identification accuracy comparison of machine learning models using odorant structure features.

| Model | Features | AUC | Precision | Recall | F1 score |
| --- | --- | --- | --- | --- | --- |
| GLVQ | Alexnet | 0.704±0.004 | 0.560±0.002 | 0.645±0.003 | 0.507±0.007 |
|  | Densenet | 0.717±0.005 | 0.565±0.002 | 0.663±0.003 | 0.510±0.008 |
|  | Restnet | 0.742±0.006 | 0.580±0.003 | 0.691±0.005 | 0.548±0.009 |
|  | VGG | 0.691±0.005 | 0.552±0.001 | 0.639±0.003 | 0.488±0.005 |
|  | Molecular parameters | 0.492±0.002 | 0.497±0.000 | 0.494±0.001 | 0.408±0.001 |
|  | Finger prints | 0.521±0.003 | 0.506±0.000 | 0.516±0.001 | 0.408±0.007 |
| GDBT | Alexnet | 0.788±0.005 | 0.589±0.004 | 0.706±0.004 | 0.562±0.007 |
|  | Densenet | 0.800±0.004 | 0.595±0.004 | 0.721±0.003 | 0.570±0.007 |
|  | Restnet | 0.790±0.004 | 0.591±0.004 | 0.719±0.004 | 0.563±0.007 |
|  | VGG | 0.778±0.005 | 0.585±0.003 | 0.710±0.004 | 0.558±0.005 |
|  | Molecular parameters | 0.538±0.003 | 0.512±0.000 | 0.530±0.002 | 0.413±0.002 |
|  | Finger prints | 0.544±0.002 | 0.512±0.000 | 0.530±0.001 | 0.421±0.002 |

**Fig. 6.**
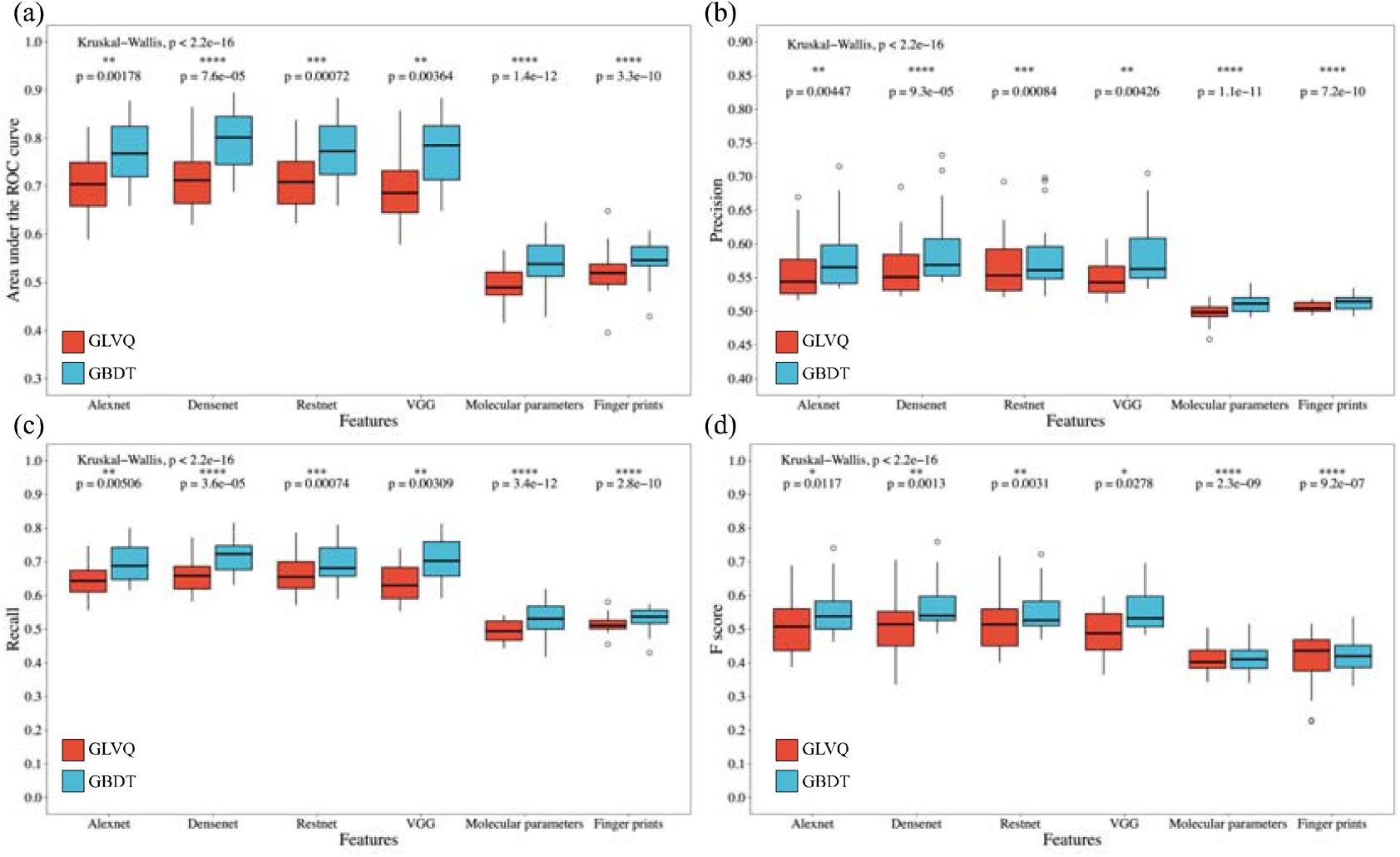
Comparison of average identification area under the ROC (a), precision (b), recall (c), and F-score (d) for the GLVQ and GBDT models under molecular graphic feature extractions, molecular parameters, and molecular fingerprints. Results were evaluated using the nonparametric Wilcoxon signed-rank test.

For the GBDT models, the parameters including the learning rate, max depth of trees, the fraction of features for each tree, gamma, min child weight, and subsample values were set to 0.2, 4, 0.5, 2, 0.5, and 0.5, respectively. As illustrated in Fig. S9, the features extracted using GBDT with VGG showed a higher prediction accuracy for cluster-1, cluster-5, and cluster-17 than for the other clusters. In addition, Restnet did a better job of identifying cluster-7, cluster-8, cluster-10, and cluster-18. However, Densenet showed better prediction performance for most clusters. The average odor cluster identification results calibrated using the GBDT models are shown in Fig. 6 and Table 2, which shows that the identification accuracy of Densenet (AUC 0.800±0.004, precision 0.595±0.004, recall 0.721±0.003, and F-score 0.570±0.007) was significantly higher than that of the other molecular feature extraction datasets (p<0.001).

### 3.7 SOR model comparison

When we compared the two modeling methods, we found that the GBDT had better identification accuracy than the GLVQ. In general, the GBDT run with features extracted via Desenet had the best identification performance (AUC 0.800±0.004, precision 0.595±0.004, recall 0.721±0.003, and F-score 0.570±0.007, p<0.001), followed by the Restnet-GBDT (AUC 0.790±0.004, precision 0.591±0.004, recall 0.719±0.004, and F-score 0.563±0.007, p<0.001) and Alexnet-GBDT (AUC 0.788±0.005, precision 0.589±0.004, recall 0.706±0.004, and F-score 0.562±0.007, p<0.001). Models trained using features extracted from molecular structure images showed higher identification accuracy (AUC 0.756±0.005, precision 0.580±0.003, recall 0.688±0.004, and F-score 0.544±0.008, p<0.001) than that calibrated using molecular parameter features (AUC 0.522±0.003, precision 0.506±0.002, recall 0.512±0.002, and F-score 0.407±0.004, p<0.001). This indicates that molecular spatial structure is more highly correlated with odor perception than pure molecular parameters, which has been previously confirmed by biology experiments [57]. A similar conclusion has been noted in related work [25]. In summary, we suggest that the GBDT run with features extracted by Densenet from molecular structure images is the optimal model for identifying perception clusters of odorants.

### 3.8 Discussion

This paper reported an odor descriptors clustering approach aimed at testing the feasibility of defining odor categories based on the co-occurrence between odor perceptions. We employed the BCE for describing co-occurrence relations between ODs from three odor databases. Results indicated that 256 ODs were clustered into 20 categories by HCA. Furthermore, proposed odor categories were supported by not only semantic analysis, but also the SOR. Marcelo et. al. reviewed the extant biological knowledge on olfaction to clarify the dimensionality of smell [58]. They suggested that although the human nose has over 400 olfactory receptors, the dimensionality of odor perception would be around 20 or less. Therefore, 20 odor categories proposed in the present study would be reasonable to understand the biology olfaction perception space.

For developing ML-GCO, we can train a model for odor category identification instead of OD identification, which would be easier to train. In the future, a reliable enough model would be developed as an odor perception recommendation system for the assessors of GC/O to reduce their burdens.

While our approach yielded OD clustering results based on the co-occurrence relationships, several limitations of the proposed approach should be discussed. First, abundant ODs, such as sweet and fruity, were not considered because of their ambiguities. Therefore, we need to define a metric to remove these ambiguous ODs and to identify characteristic ODs, such as coffee and vanilla. Second, the SOR model should be extended to include mixtures of odorants. Molecular structures cannot be arbitrarily blended by naive linear superposition. Thus, novel approaches and algorithms, such as molecular 3-dimensional interaction embedding, topology graph representations, and mixed MS analysis should be considered for use in extracting critical features for odor mixture presentation. We hope that, with these possible directions, our work will provide a foundation for understanding human olfaction space and finding the basis of odor perception.

## 4. Conclusion

In this paper, we describe a method for extracting generalized smell perceptions, termed odor categories. For clustering analysis, a metric should be defined to represent the relationship between these perceptions. In this study, we introduced BCE as a vector embedder for describing the co-occurrence relationships between ODs. The ODs of odorants were also calibrated based on the above-mentioned embedded vectors to reduce the diversity evaluation criterion in the databases. After removing infrequent ODs, cluster analyses were performed on the embedded vectors of the ODs. The results indicated that the ODs were clustered in 20 groups, and most of the ODs with similar semantic meanings were clustered in the same class.

To verify the rationality of the proposed clustering scenario, we used molecular structure features to predict the aforementioned odor categories. The high prediction accuracy (AUC is 0.8) of the SOR models demonstrated that co-occurrence-based embedded vectors may be feasible descriptors for expressing the similarity between ODs. In addition, our data demonstrate that molecular structures combined with ML methods can be adopted for odor perception cluster identification, which would be an intelligent auxiliary system for existing GC/O.

## CRediT authorship contribution statement

**Liang Shang**: Experiments, Data curation, Writing - original draft. **Chunjun Liu**: Supervision, Project integration, Writing - original draft. **Fengzhen Tang**: Supervision, Project integration. **Bin Chen**: Experiments, Data curation. **Lianqing Liu**: Supervision. **Kenshi Hayashi**: Supervision.

## Acknowledgments

This research was supported by a grant of China Postdoctoral Science Foundation (No. 2021M703399), National Key Research and Development Program of China (No. 2020YFB13400), National Nature Science Foundation of China (No. 61803369 and 61801400), and JSPS KAKENHI Grant (No. 18H03782).

## Appendix A. Supplementary data

Supplementary data to this article can be found online at http://doi.org/xxx. Detail description of algorithms and models, includes t-SNE, transductive bagging learning, model evaluation metrics, GLVQ, GDBT, supplementary figures, and tables for detailed experiment and analysis results of this work (PDF).

