## Supplementary material for "Machine-Learning-Based Olfactometry: An Auxiliary System for Human Assessors in Olfactory Measurement": SI

Table S3. Correlations of Word2Vec embedded vectors for odor categories. . 48
Table S4. Odor cluster identification results by GLVQ and GBDT models. . 49

**Reference** **55**

### Supplementary Methods

#### Barnes-Hut t-distributed Stochastic Neighbor Embedding (t-SNE)

As an unsupervised embedding algorithm, t-SNE has been widely used to visualize high-dimensional data at a lower dimension to enable dataset embeddings to be learned. The computational process was introduced as follows. Let $\mathfrak{D}$ be a data set $\mathfrak{D} = \{x_i | x_i \in R^n, i = 1, 2, \ldots, m\}$, where $x_i$ is a $n$ dimension vector of the $i$-th sample, and n is the total number of samples in the dataset. In the present study, m and n were 265 and 64, respectively. The distance between $i$ and $j$ was defined by $dist(x_i, x_j)$. We employed Euclidean distance as the distance function in this study, $dist(x_i, x_j) = |x_i - x_j|^2$.

Firstly, the pairwise similarities ($p_{ij}$) between the samples could be calculated as:

$$p_{ij} = \frac{\exp(-dist(x_i, x_j)^2/2\sigma^2)}{\sum_{k \neq l}(\exp(-dist(x_k, x_l)^2/2\sigma^2))} \tag{1}$$

where $\sigma$ was the variance parameter of the Gaussian function.

Second, the embedded target presentation ($\mathfrak{T}$) was defined as $\mathfrak{T} = \{y_i | y_i \in R^d, i = 1, 2, \ldots, m\}$, where $d$ was the dimension of the target ($d = 2$ or 3) using a normalized Student's-t kernel with one degree of freedom. The similarity $q_{ij}$ between the target values could be defined as follows.

$$q_{ij} = \frac{(1 + dist(y_i, y_j)^2)^{-1}}{\sum_{k \neq l}(1 + dist(y_k, y_l)^2)^{-1}} \tag{2}$$

Finally, the locations of the embedding points ($Y$) could be determined by Kullback-Leibler (KL) divergence. Therefore, the optimal low-dimensional representation $\mathfrak{T}$ was obtained by minimizing the $cost(\mathfrak{T})$:

$$\begin{aligned} Cost(\mathfrak{T}) &= KL(p||q) \\ &= \sum_{i \neq j} p_{ij} \log \frac{p_{ij}}{q_{ij}} \end{aligned} \tag{3}$$

A detailed explanation of t-SNE was presented elsewhere.[1]

#### Transductive Bagging Learning

The dataset for the OD cluster prediction was a typical imbalanced data set because the class distribution of the positive samples (minor samples with specified OD clusters) and negative samples (major samples with non-specified OD clusters) was not uniform. Inspired by Mordelet's research, we considered bagging classifiers to be a feasible method for learning with an imbalanced data set.[2] As described in Algorithm 1, a random subsample $N_t$ is first generated from a negative sample set ($N$). Afterwards, a classifier is calibrated to discriminate all of the positive samples ($P$) from $N_t$. The number of bootstraps ($T$) was assigned to define the boosting model number. In the present study, the sample pool was divided into training and test sets with a 3:1 ratio via random stratified sampling. In addition, the number of bootstraps ($T$) was set as 100 and the subsample number ($K$) was the same as that for the positive numbers ($P$). Considering the sample size, we employed GBDT[3] and GLVQ to predict the odor clusters in present study. Finally, the optimal feature extractor and model combination was determined by considering the area under the ROC curve (AUC-ROC), precision, recall, and F1-score of the test set, respectively.

**Algorithm 1:** Transductive bagging learning

**Input:** $P$, $N$, $K$=size of bootstrap samples, $T$=number of bootstraps

**Output:** a score s: $N \rightarrow \mathbb{R}$

```
Initialize ∀x ∈ N, n(x) ← 0, f(x) ← 0
for t=1 to T do
    Draw a bootstrap sample N_t of size K in N.
    Train a classifier f_t to discriminate P against N_t.
    for x ∈ N\N_t do
        Update:
        f(x) ← f(x) + f_t(x),
        n(x) ← n(x) + 1
Return s(x) = f(x)/n(x) for x ∈ N.
```

#### Model Evaluation Metrics

Metrics employed in the present study included the area under the ROC curve (AUC), precision, recall, and F1-score. These metrics can be defined as follows:

$$\begin{aligned} precision &= \frac{TP}{TP+FP} \\ recall &= \frac{TP}{TP+FN} \\ F1\ score &= 2 \times \frac{precision \times recall}{precision + recall} \end{aligned} \tag{4}$$

where $TP$, $FP$, and $FN$ indicate the numbers of true positives, false positives, and false negatives, respectively.

#### Generalized Learning Vector Quantization

Generalized learning vector quantization (GLVQ) is a simple and efficient method that has been widely applied in many classification scenarios, especially for electroencephalogram

(EEG) data classification.[4] As a metric comparisons-based supervised algorithm, we considered GLVQ to be suitable for exploring the internal relationships between smell perception descriptors.

Let $\mathfrak{D}$ be a data set $(\mathbf{x_i}, y_i) \in \mathbb{R}^n \times \{1, \ldots, C\}$, $i = 1, \ldots, m$ for training, where $n$ is the dimension of input data and $m$ is the number of samples. In our particular case, we needed to predict whether or not a sample belonged to a target cluster. Therefore, the number of classes $C$ was 2. Briefly, the goal of GLVQ was to automatically obtain prototypes based on the steepest descent method in a well-defined cost function. In summary, the cost function of GLVQ can be defined as follows:

$$\begin{aligned} E(\mathbf{w}) &= \sum_{i=1}^{m} \Phi(f(\mathbf{x}_i, \mathbf{w})) \\ f(\mathbf{x}_i, \mathbf{w}) &= \frac{dist(\mathbf{x}_i, \mathbf{w}_J) - dist(\mathbf{x}_i, \mathbf{w}_K)}{dist(\mathbf{x}_i, \mathbf{w}_J) + dist(\mathbf{x}_i, \mathbf{w}_K)} \end{aligned} \tag{5}$$

where $\Phi(\cdot)$ indicates an activation function and $dist(\cdot, \cdot)$ is a function that calculates the distance between 2 vectors. Here, we applied the squared Euclidean distance, i.e., $dist(\mathbf{x}_i, \mathbf{w}_J) = |\mathbf{x}_i - \mathbf{w}_J|^2$, to evaluate the differences between samples. $\mathbf{w}_J$ and $\mathbf{w}_K$ are the closest correct and incorrect prototypes to $\mathbf{w}_i$, respectively. By minimizing the cost function based on the stochastic steepest gradient descent algorithm, prototypes from GLVQ could be learned and updated. The learning rules can be defined as follows:

$$\begin{aligned} \Delta\mathbf{w}_J &= \eta(t) \frac{\partial \Phi}{\partial f} \frac{4 \times dist(\mathbf{x}_i, \mathbf{w}_K)}{(dist(\mathbf{x}_i, \mathbf{w}_J) + dist(\mathbf{x}_i, \mathbf{w}_K))^2} (\mathbf{x}_i - \mathbf{w}_J) \\ \Delta\mathbf{w}_K &= -\eta(t) \frac{\partial \Phi}{\partial f} \frac{4 \times dist(\mathbf{x}_i, \mathbf{w}_J)}{(dist(\mathbf{x}_i, \mathbf{w}_J) + dist(\mathbf{x}_i, \mathbf{w}_K))^2} (\mathbf{x}_i - \mathbf{w}_K) \end{aligned} \tag{6}$$

where learning rate ($\eta(t) \in (0, 1)$) is satisfied $\sum_{t=1}^{\infty} \eta(t) = \infty$ and $\sum_{t=1}^{\infty} \eta(t)^2 < \infty$. By updating the prototypes according to the learning rate and cost function, the GLVQ models could be learned and calibrated.

#### Gradient Boosting Decision Tree

As a powerful machine-learning technique, the gradient boosting decision tree (GBDT) has been applied in a wide range of areas, including medical and commercial settings.[5] The main idea behind GBDT is the use of binary classification to distinguish between two classes. Similarly, let $\mathfrak{D}$ be a training data set $(\mathbf{x_i}, y_i) \in \mathbb{R}^n \times \{0, 1\}$, $i = 1, \ldots, m$, where $n$ is the input dimension and $m$ is the number of samples. The goal of GBDT is to choose a classification function $F(\mathbf{x}_i)$ to minimize the aggregation of cost function $L(y_i, F(\mathbf{x}_i))$, which can be defined by:

$$\begin{aligned} F^* &= \arg\min \sum_{i=1}^{m} L(y_i, F(\mathbf{x}_i)) \\ F(\mathbf{x}) &= \sum_{j=1}^{T} f_j(\mathbf{x}) \end{aligned} \tag{7}$$

where $T$ indicates the number of iterations. $f_j(\mathbf{x})$ are designed in an incremental fashion. Specifically, at the $j$-th stage, the added function $(f_j)$ is selected to optimize the aggregated loss while keeping $\{f_k\}_{k=1}^{j-1}\}$ fixed. Each function $f_j$ belongs to a set of parametrized decision trees. Let $\theta$ denote the parameters of the decision trees, such as the number of features to split for each node, etc. For the $j$-th stage, the cost function can be defined as follows.

$$\begin{aligned} L(y_i, F_{j-1}(\mathbf{x}_i) + f_m(\mathbf{x}_i)) \approx \\ L(y_i, F_{j-1}(\mathbf{x}_i)) + \nabla_i f_m(\mathbf{x}_i) + \frac{1}{2} f_m(\mathbf{x}_i)^2 \\ \nabla_i = \frac{\partial L(y_i, F(\mathbf{x}_i))}{\partial F(\mathbf{x}_i)} |_{F(\mathbf{x}_i)=F_{j-1}(\mathbf{x}_i)} \end{aligned} \tag{8}$$

By selecting $f_j$ to minimize the right part of the cost function, the optimization function can be written as the following minimization problem:

$$\arg\min_{f_j} \sum_{i=1}^{m} \frac{1}{2} (f_m(\mathbf{x}_i) - \nabla_i)^2 \tag{9}$$

### Supplementary Figures and Tables

#### Supplementary Figures

- Figure S1. BCE embedding dimension selection based on the normalized discounted cumulative gain of the top 20 samples (NDCG@20).
- Figure S2. Distribution of odor descriptor sample size before and after relabeling by the BCE algorithm.
- Figure S3.Odor descriptor cluster analysis based on the co-occurrence of embedded vectors.
- Figure S4. Hierarchical clustering based on odor descriptors embedded vectors from BCE method.
- Figure S5. Correlation heat-maps for pre-trained Word2Vec embedded vectors for smell categories.
- Figure S6. Distribution plots of correlation between Word2Vec embedded odor descriptor vectors.
- Figure S7. Sample distribution for each odor cluster and Statistical distribution of the number of clusters for odorants.
- Figure S8. Comparison of odor clusters identification performance by GLVQ models.
- Figure S9. Comparison of odor clusters identification performance by GDBT models

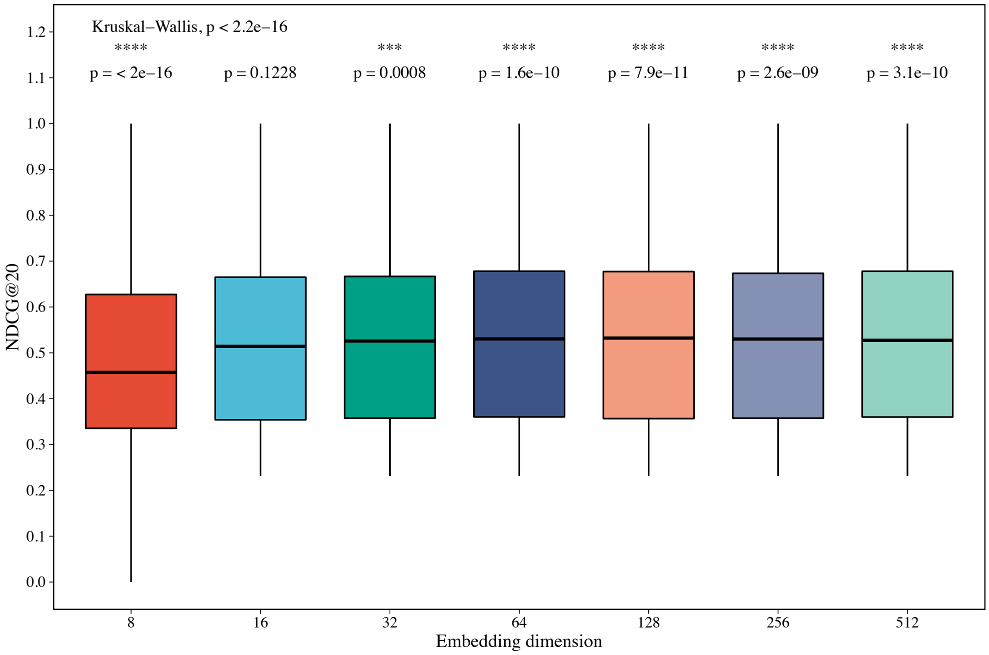

Figure S1: BCE embedding dimension selection based on the normalized discounted cumulative gain of the top 20 samples (NDCG@20).

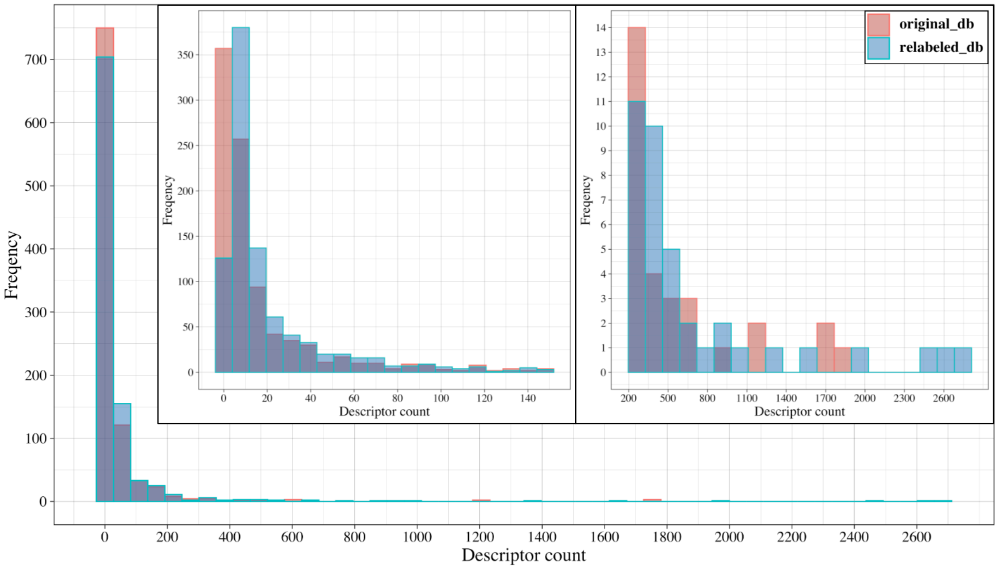

Figure S2: Distribution of odor descriptor sample size before and after relabeling by the BCE algorithm.

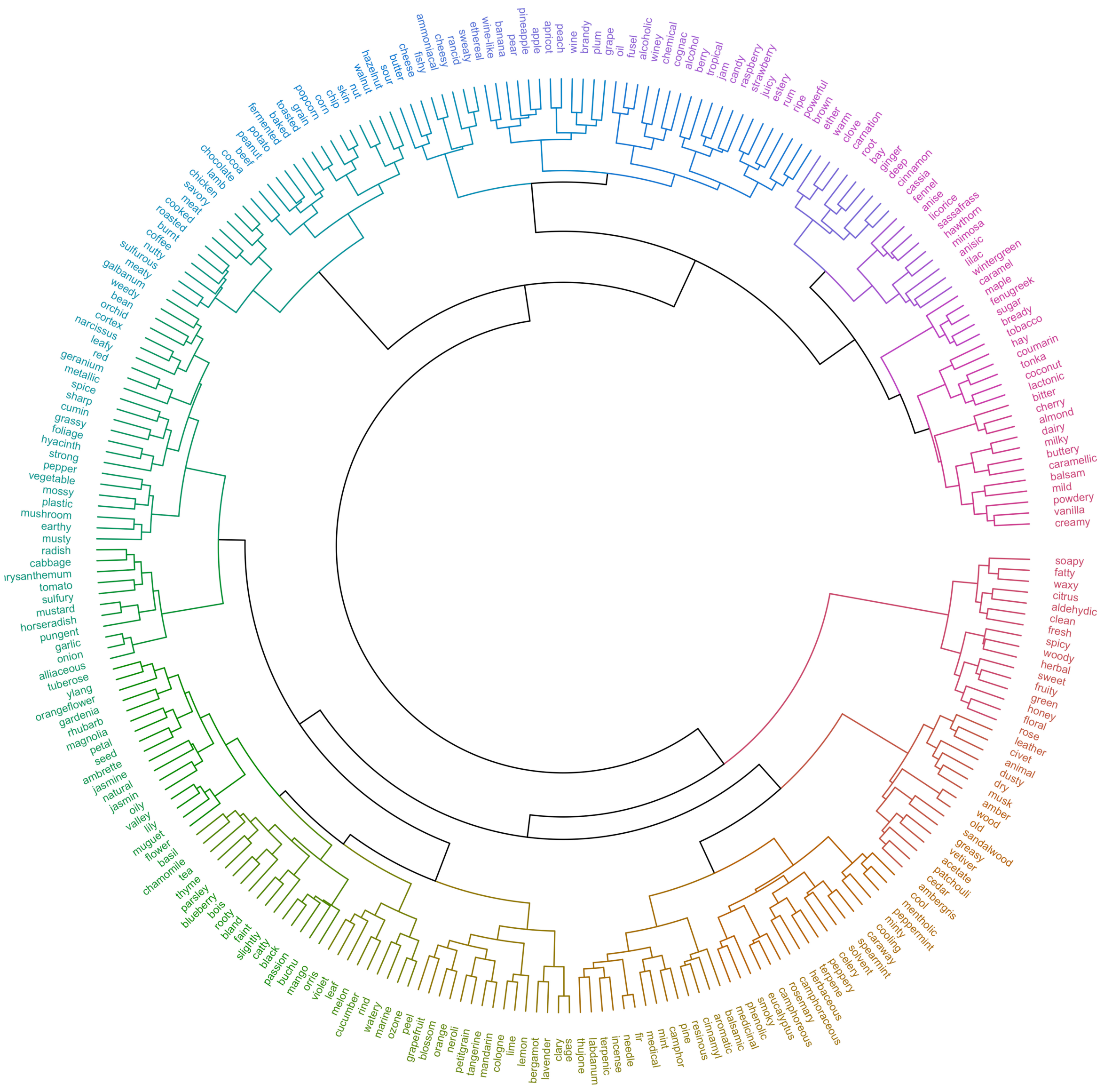

Figure S3: Odor descriptor cluster analysis based on the co-occurrence of embedded vectors.

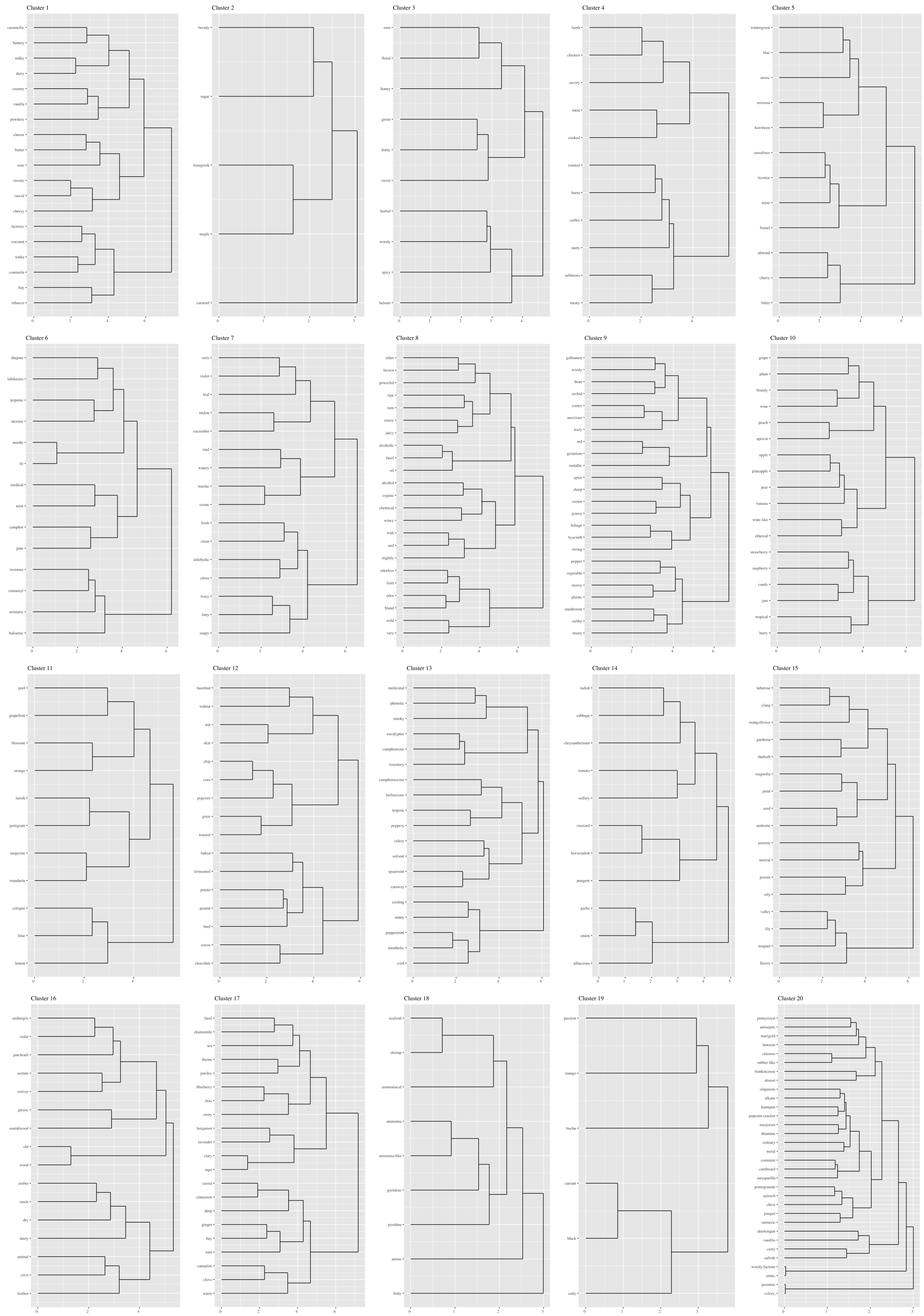

Figure S4: Hierarchical clustering based on odor descriptors embedded vectors from BCE method.

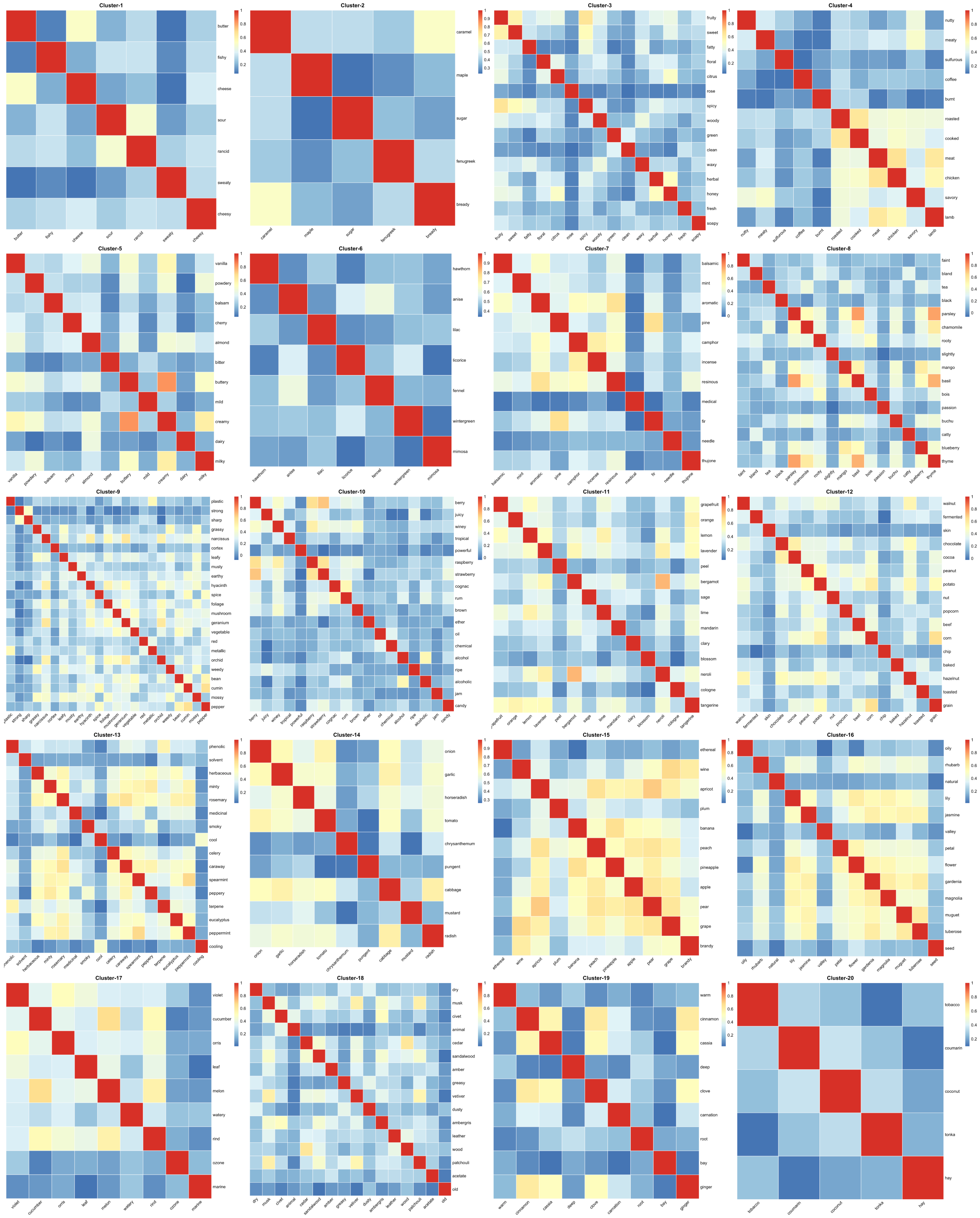

Figure S5: Correlation heat-maps for pre-trained Word2Vec embedded vectors for smell categories.

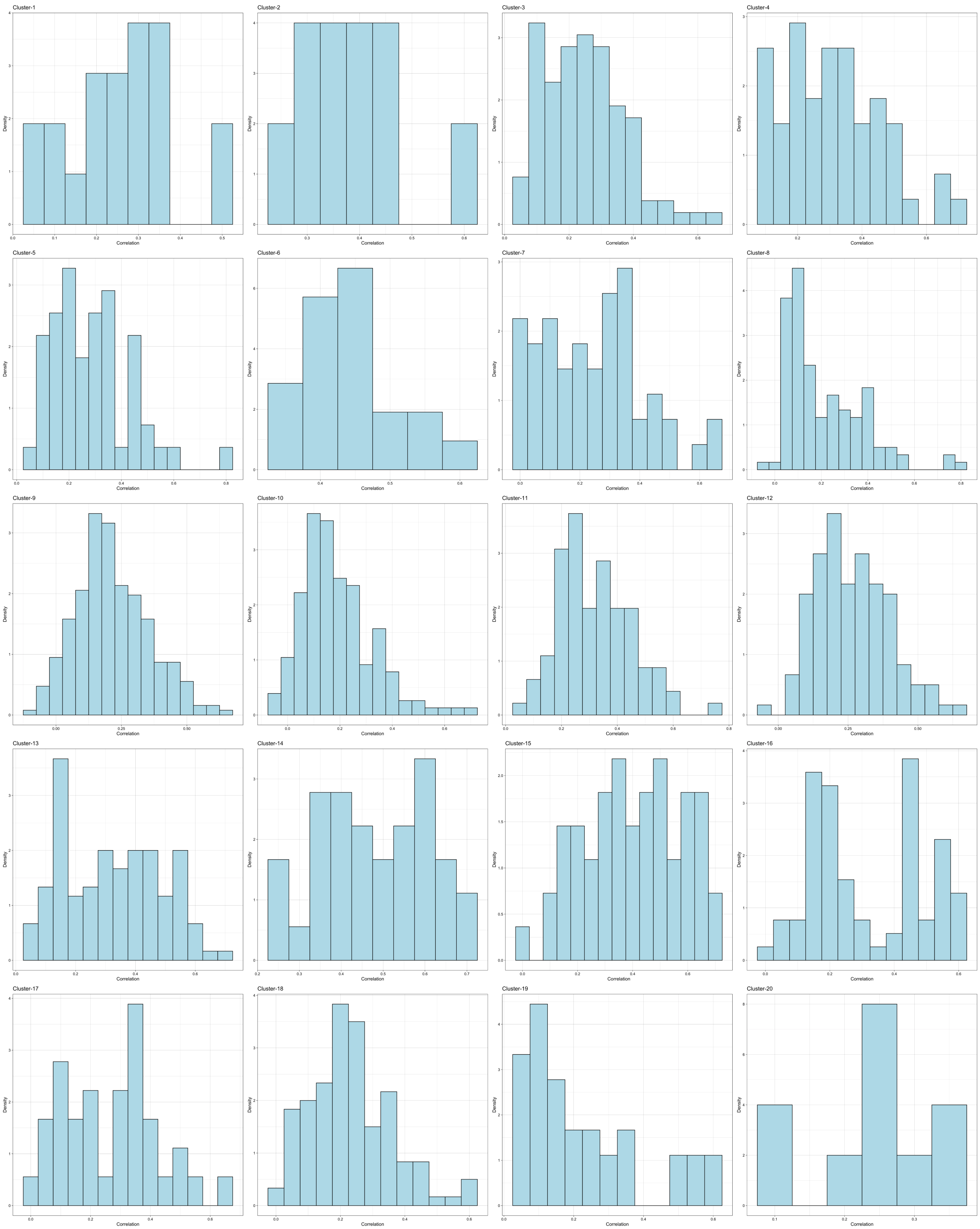

Figure S6: Distribution plots of correlation between Word2Vec embedded odor descriptor vectors.

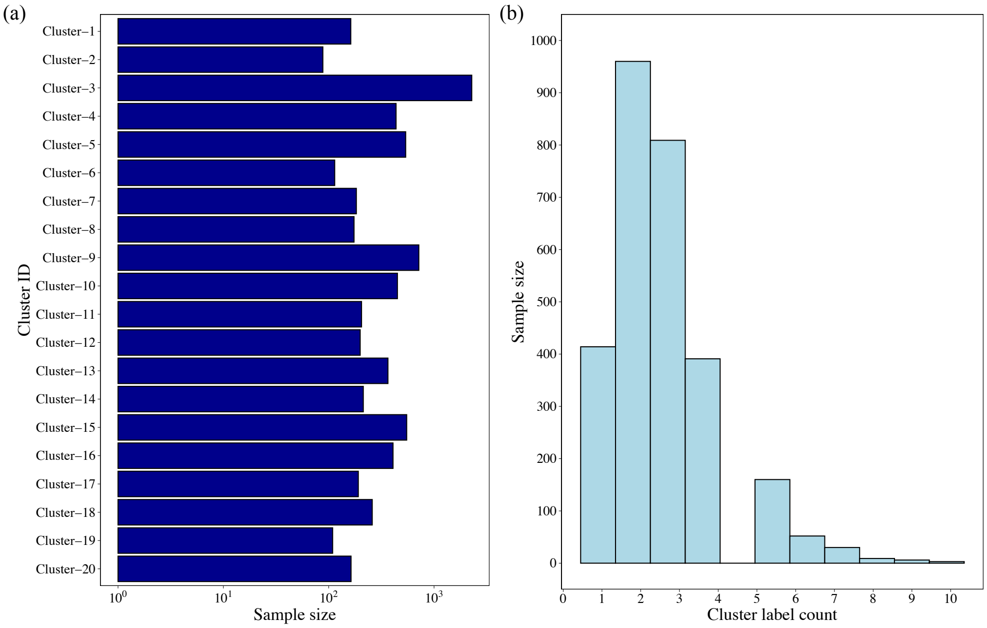

Figure S7: Sample distribution for each odor cluster (a). Statistical distribution of the number of clusters for odorants (b). Most odorants belonged to more than two clusters, which corresponded to the complexity and ambiguity of odor perceptions.

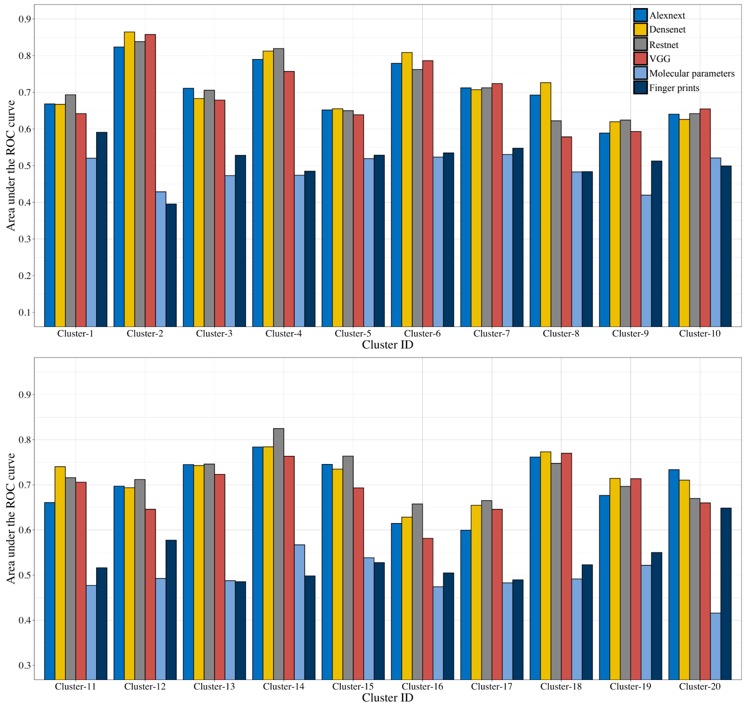

(a) Area under the ROC curve for GLVQ model

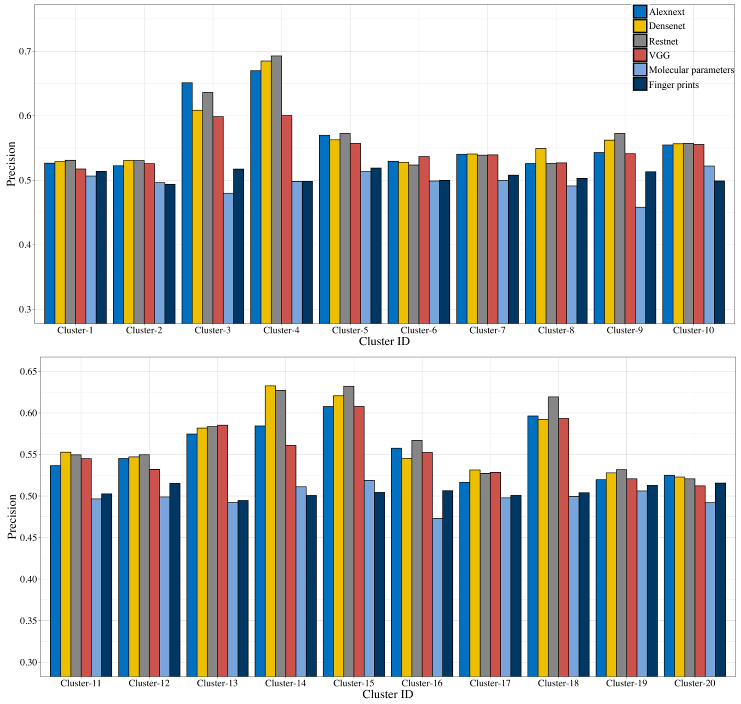

(b) Precision for GLVQ model

Figure S8: Comparison of odor clusters identification for area under the ROC (a), precision (b), recall (c) and F-score (d) by GLVQ models under CNNs feature extraction (Alexnet, Densenet, Restnet or VGG), molecularly parameters or molecular finger prints.

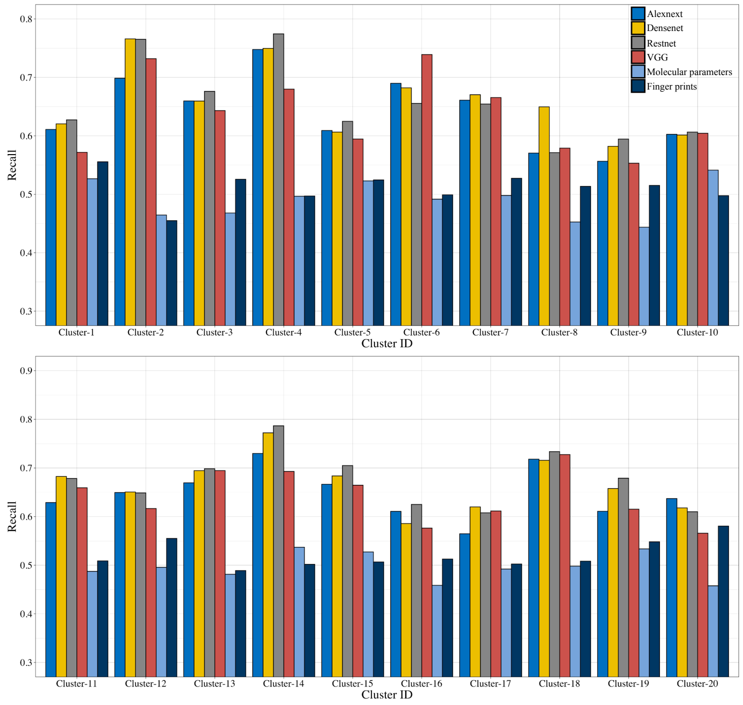

(c) Recall for GLVQ model

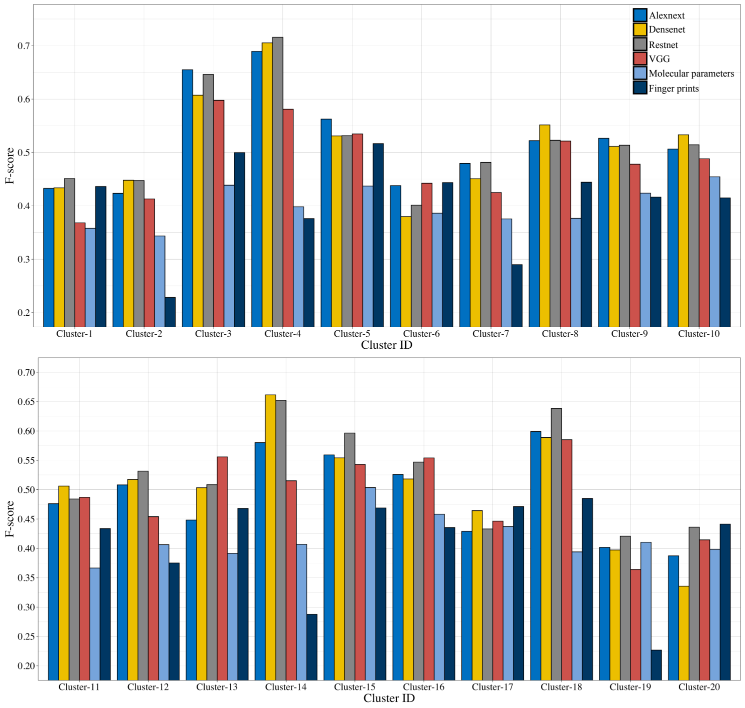

(d) F-score for GLVQ model

Figure S8: Comparison of odor clusters identification for area under the ROC (a), precision (b), recall (c) and F-score (d) by GLVQ models under CNNs feature extraction (Alexnet, Densenet, Restnet or VGG), molecularly parameters or molecular finger prints.

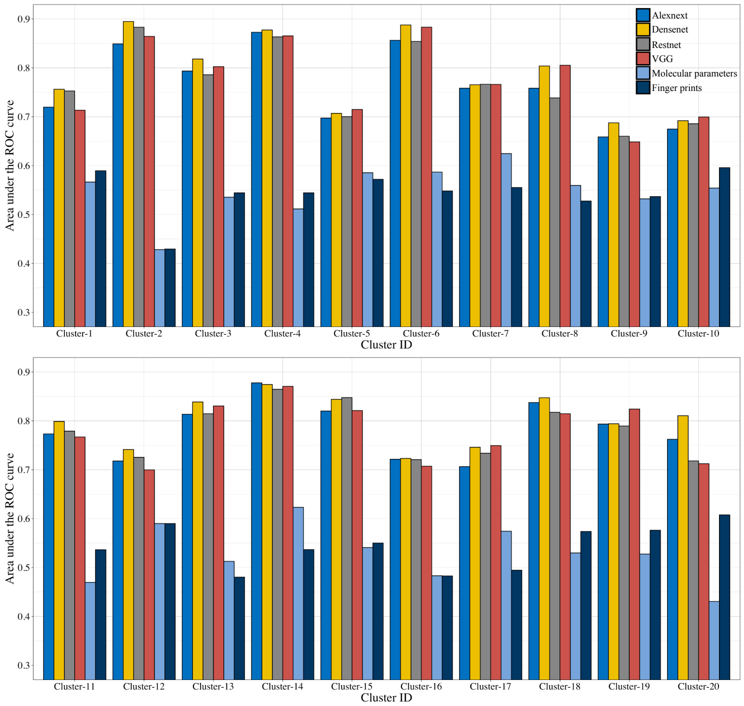

(a) Area under the ROC curve for GBDT model

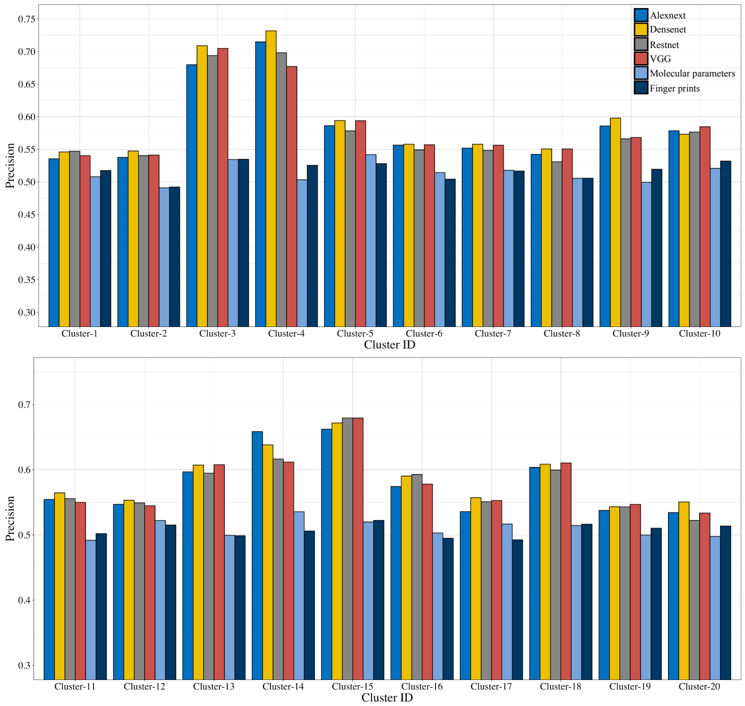

(b) Precision for GBDT model

Figure S9: Comparison of odor clusters identification for area under the ROC (a), precision (b), recall (c) and F-score (d) by GBDT models under CNNs feature extraction (Alexnet, Densenet, Restnet or VGG), molecularly parameters or molecular finger prints.

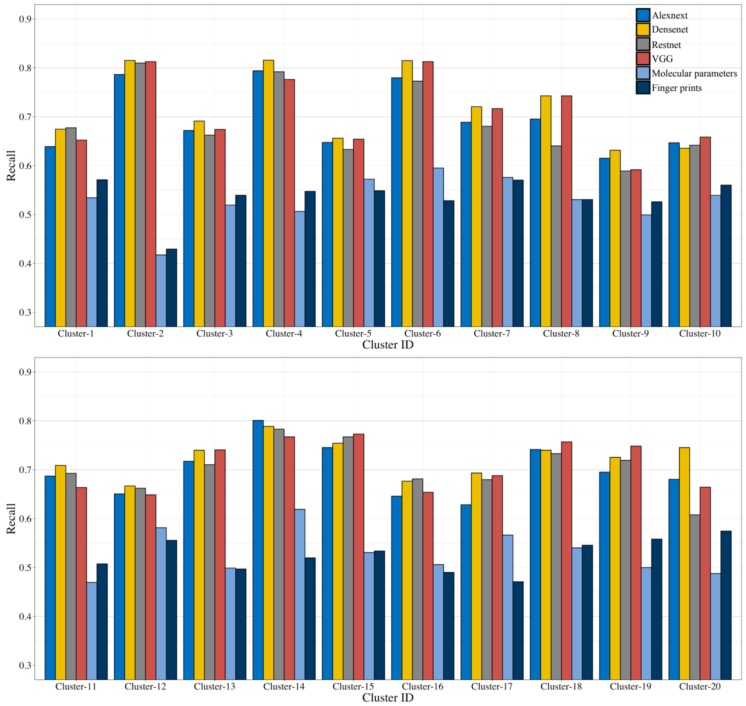

(c) Recall for GBDT model

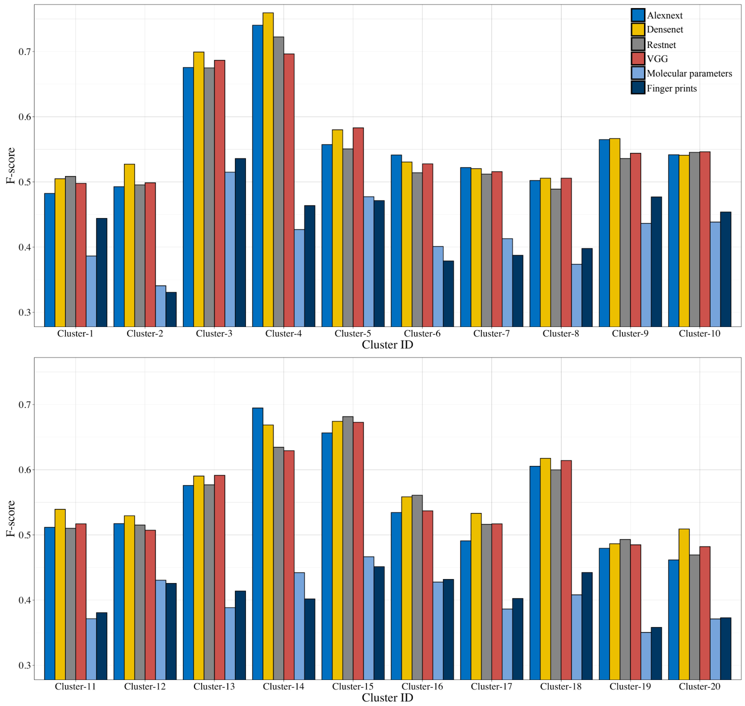

(d) F-score for GBDT model

Figure S9: Comparison of odor clusters identification for area under the ROC (a), precision (b), recall (c) and F-score (d) by GBDT models under CNNs feature extraction (Alexnet, Densenet, Restnet or VGG), molecularly parameters or molecular finger prints.

#### Supplementary Tables

- Table S1. Odor descriptors distribution before and after calibrated by BCE method.
- Table S2. Detail information for t-SNE plots and odor descriptors clustering results.
- Table S3. Correlations of Word2Vec embedded vectors for odor categories.
- Table S4. Odor clusters identification results by GLVQ and GBDT models under Alexnet (a), Densenet (b), Restnet (c), VGG (d), molecularly parameters (e) and molecular finger prints (f).

Table S1: Odor descriptors distribution before and after calibrated by BCE method.

| Descriptor | Original Count | Relabeled Count | Difference | Considered or not |
|---|---|---|---|---|
| sweet | 1758 | 2686 | 928 | TRUE |
| green | 1760 | 2648 | 888 | TRUE |
| fruity | 1780 | 2460 | 680 | TRUE |
| floral | 1200 | 1962 | 762 | TRUE |
| woody | 1184 | 1630 | 446 | TRUE |
| herbal | 904 | 1364 | 460 | TRUE |
| waxy | 582 | 982 | 400 | TRUE |
| fresh | 674 | 934 | 260 | TRUE |
| spicy | 624 | 864 | 240 | TRUE |
| fatty | 594 | 762 | 168 | TRUE |
| citrus | 486 | 656 | 170 | TRUE |
| rose | 508 | 656 | 148 | TRUE |
| sulfurous | 346 | 574 | 228 | TRUE |
| apple | 318 | 538 | 220 | TRUE |
| meaty | 278 | 512 | 234 | TRUE |
| oily | 380 | 498 | 118 | TRUE |
| nutty | 352 | 496 | 144 | TRUE |
| roasted | 240 | 434 | 194 | TRUE |
| earthy | 412 | 432 | 20 | TRUE |
| tropical | 314 | 432 | 118 | TRUE |
| pineapple | 252 | 406 | 154 | TRUE |
| onion | 158 | 372 | 214 | TRUE |
| dry | 240 | 348 | 108 | TRUE |
| creamy | 254 | 336 | 82 | TRUE |

Table S1: Odor descriptors distribution before and after calibrated by BCE method.

| Descriptor | Original Count | Relabeled Count | Difference | Considered or not |
|---|---|---|---|---|
| minty | 286 | 334 | 48 | TRUE |
| coffee | 156 | 330 | 174 | TRUE |
| vegetable | 238 | 328 | 90 | TRUE |
| balsam | 302 | 304 | 2 | TRUE |
| garlic | 114 | 272 | 158 | TRUE |
| ethereal | 240 | 260 | 20 | TRUE |
| musty | 244 | 244 | 0 | TRUE |
| powdery | 216 | 236 | 20 | TRUE |
| sulfury | 90 | 226 | 136 | TRUE |
| banana | 176 | 224 | 48 | TRUE |
| honey | 212 | 222 | 10 | TRUE |
| amber | 150 | 206 | 56 | TRUE |
| caramel | 168 | 202 | 34 | TRUE |
| pear | 174 | 202 | 28 | TRUE |
| mild | 196 | 198 | 2 | TRUE |
| balsamic | 190 | 192 | 2 | TRUE |
| phenolic | 184 | 192 | 8 | TRUE |
| camphor | 154 | 190 | 36 | TRUE |
| coconut | 172 | 190 | 18 | TRUE |
| berry | 186 | 186 | 0 | TRUE |
| aldehydic | 170 | 184 | 14 | TRUE |
| musk | 146 | 182 | 36 | TRUE |
| orange | 160 | 178 | 18 | TRUE |
| vanilla | 176 | 176 | 0 | TRUE |

Table S1: Odor descriptors distribution before and after calibrated by BCE method.

| Descriptor | Original Count | Relabeled Count | Difference | Considered or not |
|---|---|---|---|---|
| wine-like | 162 | 166 | 4 | TRUE |
| alliaceous | 70 | 164 | 94 | TRUE |
| clean | 154 | 160 | 6 | TRUE |
| melon | 146 | 160 | 14 | TRUE |
| faint | 82 | 158 | 76 | TRUE |
| tobacco | 152 | 158 | 6 | TRUE |
| pine | 116 | 156 | 40 | TRUE |
| burnt | 106 | 154 | 48 | TRUE |
| metallic | 154 | 154 | 0 | TRUE |
| animal | 126 | 152 | 26 | TRUE |
| chocolate | 136 | 148 | 12 | TRUE |
| cheese | 130 | 146 | 16 | TRUE |
| herbaceous | 146 | 146 | 0 | TRUE |
| jasmin | 144 | 144 | 0 | TRUE |
| lemon | 130 | 144 | 14 | TRUE |
| odorless | 100 | 144 | 44 | TRUE |
| peach | 138 | 144 | 6 | TRUE |
| apricot | 132 | 138 | 6 | TRUE |
| cherry | 128 | 136 | 8 | TRUE |
| almond | 120 | 130 | 10 | TRUE |
| cooked | 86 | 122 | 36 | TRUE |
| leafy | 120 | 120 | 0 | TRUE |
| mushroom | 118 | 120 | 2 | TRUE |
| violet | 116 | 120 | 4 | TRUE |

Table S1: Odor descriptors distribution before and after calibrated by BCE method.

| Descriptor | Original Count | Relabeled Count | Difference | Considered or not |
|---|---|---|---|---|
| warm | 118 | 118 | 0 | TRUE |
| cedar | 86 | 116 | 30 | TRUE |
| grape | 114 | 114 | 0 | TRUE |
| cooling | 44 | 110 | 66 | TRUE |
| anise | 94 | 106 | 12 | TRUE |
| medicinal | 96 | 106 | 10 | TRUE |
| natural | 106 | 106 | 0 | TRUE |
| geranium | 92 | 104 | 12 | TRUE |
| mentholic | 36 | 104 | 68 | TRUE |
| pungent | 100 | 104 | 4 | TRUE |
| butter | 100 | 102 | 2 | TRUE |
| cinnamon | 82 | 100 | 18 | TRUE |
| buttery | 88 | 98 | 10 | TRUE |
| cocoa | 82 | 94 | 12 | TRUE |
| dairy | 76 | 94 | 18 | TRUE |
| hay | 90 | 92 | 2 | TRUE |
| lily | 84 | 92 | 8 | TRUE |
| plum | 92 | 92 | 0 | TRUE |
| spearmint | 38 | 92 | 54 | TRUE |
| grapefruit | 90 | 90 | 0 | TRUE |
| hyacinth | 90 | 90 | 0 | TRUE |
| soapy | 90 | 90 | 0 | TRUE |
| clove | 74 | 88 | 14 | TRUE |
| smoky | 68 | 88 | 20 | TRUE |

Table S1: Odor descriptors distribution before and after calibrated by BCE method.

| Descriptor | Original Count | Relabeled Count | Difference | Considered or not |
|---|---|---|---|---|
| spice | 88 | 88 | 0 | TRUE |
| cucumber | 82 | 86 | 4 | TRUE |
| lime | 64 | 84 | 20 | TRUE |
| camphoreous | 64 | 82 | 18 | TRUE |
| fishy | 72 | 82 | 10 | TRUE |
| fruit | 80 | 80 | 0 | TRUE |
| strawberry | 80 | 80 | 0 | TRUE |
| cabbage | 42 | 78 | 36 | TRUE |
| cortex | 74 | 76 | 2 | TRUE |
| maple | 50 | 76 | 26 | TRUE |
| raspberry | 76 | 76 | 0 | TRUE |
| orris | 74 | 74 | 0 | TRUE |
| bland | 22 | 72 | 50 | TRUE |
| licorice | 54 | 72 | 18 | TRUE |
| rosemary | 36 | 72 | 36 | TRUE |
| weedy | 72 | 72 | 0 | TRUE |
| tea | 70 | 70 | 0 | TRUE |
| bergamot | 50 | 68 | 18 | TRUE |
| grassy | 68 | 68 | 0 | TRUE |
| lavender | 60 | 68 | 8 | TRUE |
| meat | 46 | 68 | 22 | TRUE |
| pepper | 68 | 68 | 0 | TRUE |
| peppermint | 36 | 68 | 32 | TRUE |
| caramellic | 60 | 66 | 6 | TRUE |

Table S1: Odor descriptors distribution before and after calibrated by BCE method.

| Descriptor | Original Count | Relabeled Count | Difference | Considered or not |
|---|---|---|---|---|
| hazelnut | 64 | 66 | 2 | TRUE |
| coumarin | 58 | 64 | 6 | TRUE |
| leaf | 64 | 64 | 0 | TRUE |
| skin | 64 | 64 | 0 | TRUE |
| horseradish | 22 | 62 | 40 | TRUE |
| mint | 60 | 62 | 2 | TRUE |
| sharp | 62 | 62 | 0 | TRUE |
| wine | 62 | 62 | 0 | TRUE |
| milky | 56 | 60 | 4 | TRUE |
| savory | 42 | 60 | 18 | TRUE |
| celery | 58 | 58 | 0 | TRUE |
| nut | 52 | 58 | 6 | TRUE |
| sandalwood | 56 | 58 | 2 | TRUE |
| strong | 58 | 58 | 0 | TRUE |
| winey | 54 | 58 | 4 | TRUE |
| black | 54 | 56 | 2 | TRUE |
| gardenia | 56 | 56 | 0 | TRUE |
| muguet | 52 | 56 | 4 | TRUE |
| rum | 56 | 56 | 0 | TRUE |
| sour | 56 | 56 | 0 | TRUE |
| corn | 32 | 54 | 22 | TRUE |
| peppery | 54 | 54 | 0 | TRUE |
| powerful | 54 | 54 | 0 | TRUE |
| sugar | 28 | 54 | 26 | TRUE |

Table S1: Odor descriptors distribution before and after calibrated by BCE method.

| Descriptor | Original Count | Relabeled Count | Difference | Considered or not |
|---|---|---|---|---|
| vetiver | 50 | 54 | 4 | TRUE |
| bitter | 50 | 52 | 2 | TRUE |
| leather | 52 | 52 | 0 | TRUE |
| ripe | 52 | 52 | 0 | TRUE |
| ambergris | 36 | 50 | 14 | TRUE |
| cumin | 50 | 50 | 0 | TRUE |
| lactonic | 44 | 50 | 6 | TRUE |
| peanut | 40 | 50 | 10 | TRUE |
| potato | 46 | 50 | 4 | TRUE |
| currant | 42 | 48 | 6 | TRUE |
| tonka | 46 | 48 | 2 | TRUE |
| eucalyptus | 28 | 46 | 18 | TRUE |
| hawthorn | 42 | 46 | 4 | TRUE |
| narcissus | 46 | 46 | 0 | TRUE |
| resinous | 42 | 46 | 4 | TRUE |
| mandarin | 38 | 44 | 6 | TRUE |
| very | 34 | 44 | 10 | TRUE |
| aromatic | 42 | 42 | 0 | TRUE |
| brown | 36 | 42 | 6 | TRUE |
| ozone | 42 | 42 | 0 | TRUE |
| peel | 42 | 42 | 0 | TRUE |
| red | 36 | 42 | 6 | TRUE |
| terpene | 42 | 42 | 0 | TRUE |
| anisic | 40 | 40 | 0 | TRUE |

Table S1: Odor descriptors distribution before and after calibrated by BCE method.

| Descriptor | Original Count | Relabeled Count | Difference | Considered or not |
|---|---|---|---|---|
| cognac | 40 | 40 | 0 | TRUE |
| fir | 32 | 40 | 8 | TRUE |
| sage | 36 | 40 | 4 | TRUE |
| tomato | 40 | 40 | 0 | TRUE |
| wood | 38 | 40 | 2 | TRUE |
| ammoniacal | 24 | 38 | 14 | TRUE |
| blossom | 28 | 38 | 10 | TRUE |
| jasmine | 38 | 38 | 0 | TRUE |
| juicy | 38 | 38 | 0 | TRUE |
| mimosa | 32 | 38 | 6 | TRUE |
| neroli | 30 | 38 | 8 | TRUE |
| patchouli | 34 | 38 | 4 | TRUE |
| terpenic | 38 | 38 | 0 | TRUE |
| beef | 30 | 36 | 6 | TRUE |
| caraway | 30 | 36 | 6 | TRUE |
| fennel | 36 | 36 | 0 | TRUE |
| needle | 30 | 36 | 6 | TRUE |
| oil | 34 | 36 | 2 | TRUE |
| orangeflower | 36 | 36 | 0 | TRUE |
| plastic | 36 | 36 | 0 | TRUE |
| sassafrass | 12 | 36 | 24 | TRUE |
| toasted | 30 | 36 | 6 | TRUE |
| watery | 36 | 36 | 0 | TRUE |
| alcoholic | 30 | 34 | 4 | TRUE |

Table S1: Odor descriptors distribution before and after calibrated by BCE method.

| Descriptor | Original Count | Relabeled Count | Difference | Considered or not |
|---|---|---|---|---|
| carnation | 26 | 34 | 8 | TRUE |
| cassia | 14 | 34 | 20 | TRUE |
| cologne | 20 | 34 | 14 | TRUE |
| fusel | 28 | 34 | 6 | TRUE |
| galbanum | 34 | 34 | 0 | TRUE |
| old | 18 | 34 | 16 | TRUE |
| cheesy | 32 | 32 | 0 | TRUE |
| chip | 18 | 32 | 14 | TRUE |
| dusty | 32 | 32 | 0 | TRUE |
| jam | 32 | 32 | 0 | TRUE |
| lilac | 32 | 32 | 0 | TRUE |
| mustard | 18 | 32 | 14 | TRUE |
| orchid | 32 | 32 | 0 | TRUE |
| petal | 30 | 32 | 2 | TRUE |
| popcorn | 28 | 32 | 4 | TRUE |
| alcohol | 30 | 30 | 0 | TRUE |
| chicken | 28 | 30 | 2 | TRUE |
| clary | 26 | 30 | 4 | TRUE |
| fermented | 26 | 30 | 4 | TRUE |
| labdanum | 24 | 30 | 6 | TRUE |
| marine | 30 | 30 | 0 | TRUE |
| rooty | 30 | 30 | 0 | TRUE |
| sweaty | 28 | 30 | 2 | TRUE |
| ylang | 30 | 30 | 0 | TRUE |

Table S1: Odor descriptors distribution before and after calibrated by BCE method.

| Descriptor | Original Count | Relabeled Count | Difference | Considered or not |
|---|---|---|---|---|
| bready | 12 | 28 | 16 | TRUE |
| camphoraceous | 28 | 28 | 0 | TRUE |
| flower | 26 | 28 | 2 | TRUE |
| greasy | 28 | 28 | 0 | TRUE |
| petitgrain | 28 | 28 | 0 | TRUE |
| radish | 22 | 28 | 6 | TRUE |
| rancid | 20 | 28 | 8 | TRUE |
| solvent | 28 | 28 | 0 | TRUE |
| bay | 12 | 26 | 14 | TRUE |
| candy | 26 | 26 | 0 | TRUE |
| catty | 14 | 26 | 12 | TRUE |
| chamomile | 26 | 26 | 0 | TRUE |
| cinnamyl | 22 | 26 | 4 | TRUE |
| cool | 16 | 26 | 10 | TRUE |
| ether | 24 | 26 | 2 | TRUE |
| foliage | 26 | 26 | 0 | TRUE |
| magnolia | 24 | 26 | 2 | TRUE |
| rhubarb | 26 | 26 | 0 | TRUE |
| rind | 20 | 26 | 6 | TRUE |
| tangerine | 16 | 26 | 10 | TRUE |
| acetate | 24 | 24 | 0 | TRUE |
| ambrette | 24 | 24 | 0 | TRUE |
| buchu | 24 | 24 | 0 | TRUE |
| mango | 24 | 24 | 0 | TRUE |

Table S1: Odor descriptors distribution before and after calibrated by BCE method.

| Descriptor | Original Count | Relabeled Count | Difference | Considered or not |
|---|---|---|---|---|
| mossy | 24 | 24 | 0 | TRUE |
| thujone | 16 | 24 | 8 | TRUE |
| thyme | 24 | 24 | 0 | TRUE |
| tuberose | 22 | 24 | 2 | TRUE |
| basil | 22 | 22 | 0 | TRUE |
| chemical | 22 | 22 | 0 | TRUE |
| fenugreek | 14 | 22 | 8 | TRUE |
| ginger | 16 | 22 | 6 | TRUE |
| grain | 14 | 22 | 8 | TRUE |
| medical | 16 | 22 | 6 | TRUE |
| parsley | 16 | 22 | 6 | TRUE |
| walnut | 22 | 22 | 0 | TRUE |
| baked | 20 | 20 | 0 | TRUE |
| bean | 20 | 20 | 0 | TRUE |
| blueberry | 20 | 20 | 0 | TRUE |
| bois | 14 | 20 | 6 | TRUE |
| brandy | 20 | 20 | 0 | TRUE |
| chrysanthemum | 20 | 20 | 0 | TRUE |
| civet | 14 | 20 | 6 | TRUE |
| deep | 20 | 20 | 0 | TRUE |
| estery | 20 | 20 | 0 | TRUE |
| incense | 20 | 20 | 0 | TRUE |
| passion | 18 | 20 | 2 | TRUE |
| root | 20 | 20 | 0 | TRUE |

Table S1: Odor descriptors distribution before and after calibrated by BCE method.

| Descriptor | Original Count | Relabeled Count | Difference | Considered or not |
|---|---|---|---|---|
| seed | 18 | 20 | 2 | TRUE |
| slightly | 20 | 20 | 0 | TRUE |
| valley | 14 | 20 | 6 | TRUE |
| wintergreen | 20 | 20 | 0 | TRUE |
| absolute | 16 | 18 | 2 | FALSE |
| acetophenone | 16 | 18 | 2 | FALSE |
| acidic | 16 | 18 | 2 | FALSE |
| bacon | 16 | 18 | 2 | FALSE |
| bread | 18 | 18 | 0 | FALSE |
| carrot | 18 | 18 | 0 | FALSE |
| citronella | 18 | 18 | 0 | FALSE |
| cotton | 14 | 18 | 4 | FALSE |
| dried | 16 | 18 | 2 | FALSE |
| fecal | 14 | 18 | 4 | FALSE |
| forest | 12 | 18 | 6 | FALSE |
| gassy | 18 | 18 | 0 | FALSE |
| grass | 18 | 18 | 0 | FALSE |
| leathery | 18 | 18 | 0 | FALSE |
| linalool | 12 | 18 | 6 | FALSE |
| milk | 18 | 18 | 0 | FALSE |
| nutmeg | 18 | 18 | 0 | FALSE |
| pea | 16 | 18 | 2 | FALSE |
| peony | 16 | 18 | 2 | FALSE |
| raw | 18 | 18 | 0 | FALSE |

Table S1: Odor descriptors distribution before and after calibrated by BCE method.

| Descriptor | Original Count | Relabeled Count | Difference | Considered or not |
|---|---|---|---|---|
| rich | 18 | 18 | 0 | FALSE |
| rue | 14 | 18 | 4 | FALSE |
| beefy | 12 | 16 | 4 | FALSE |
| blackberry | 16 | 16 | 0 | FALSE |
| cedarwood | 14 | 16 | 2 | FALSE |
| clover | 16 | 16 | 0 | FALSE |
| coriander | 12 | 16 | 4 | FALSE |
| dill | 14 | 16 | 2 | FALSE |
| fungal | 14 | 16 | 2 | FALSE |
| hawthorne | 12 | 16 | 4 | FALSE |
| indole | 14 | 16 | 2 | FALSE |
| kiwi | 14 | 16 | 2 | FALSE |
| laundered | 14 | 16 | 2 | FALSE |
| myrrh | 12 | 16 | 4 | FALSE |
| oakmoss | 16 | 16 | 0 | FALSE |
| opoponax | 12 | 16 | 4 | FALSE |
| privet | 16 | 16 | 0 | FALSE |
| soft | 16 | 16 | 0 | FALSE |
| watermelon | 14 | 16 | 2 | FALSE |
| wet | 16 | 16 | 0 | FALSE |
| acid | 14 | 14 | 0 | FALSE |
| animalic | 12 | 14 | 2 | FALSE |
| bell | 14 | 14 | 0 | FALSE |
| coumarinic | 14 | 14 | 0 | FALSE |

Table S1: Odor descriptors distribution before and after calibrated by BCE method.

| Descriptor | Original Count | Relabeled Count | Difference | Considered or not |
|---|---|---|---|---|
| cyclamen | 14 | 14 | 0 | FALSE |
| fried | 12 | 14 | 2 | FALSE |
| guaiacwood | 14 | 14 | 0 | FALSE |
| honeysuckle | 14 | 14 | 0 | FALSE |
| jammy | 12 | 14 | 2 | FALSE |
| menthol | 14 | 14 | 0 | FALSE |
| naphthyl | 12 | 14 | 2 | FALSE |
| rummy | 14 | 14 | 0 | FALSE |
| saffron | 12 | 14 | 2 | FALSE |
| sawdust | 12 | 14 | 2 | FALSE |
| seedy | 14 | 14 | 0 | FALSE |
| unripe | 14 | 14 | 0 | FALSE |
| whiskey | 14 | 14 | 0 | FALSE |
| wormwood | 12 | 14 | 2 | FALSE |
| artichoke | 12 | 12 | 0 | FALSE |
| blackcurrant | 12 | 12 | 0 | FALSE |
| cloth | 12 | 12 | 0 | FALSE |
| cypress | 12 | 12 | 0 | FALSE |
| heliotrope | 12 | 12 | 0 | FALSE |
| light | 12 | 12 | 0 | FALSE |
| papaya | 12 | 12 | 0 | FALSE |
| peely | 12 | 12 | 0 | FALSE |
| rubbery | 12 | 12 | 0 | FALSE |
| seaweed | 12 | 12 | 0 | FALSE |

Table S1: Odor descriptors distribution before and after calibrated by BCE method.

| Descriptor | Original Count | Relabeled Count | Difference | Considered or not |
|---|---|---|---|---|
| tallow | 12 | 12 | 0 | FALSE |
| truffle | 12 | 12 | 0 | FALSE |

Table S2: Detail information for t-SNE plots and odor descriptors clustering results.

| Cluster ID | Descriptor | tSNE-1 | tSNE-2 |
|---|---|---|---|
| Cluster-1 | butter | -8.63280366 | -65.96061615 |
| Cluster-1 | fishy | -27.97407608 | -57.78554179 |
| Cluster-1 | ammoniacal | -28.3970502 | -56.60020128 |
| Cluster-1 | cheese | -10.0596208 | -66.83626524 |
| Cluster-1 | sour | -12.22320591 | -67.77061597 |
| Cluster-1 | rancid | -16.67077901 | -65.63607001 |
| Cluster-1 | sweaty | -17.98747806 | -65.52151726 |
| Cluster-1 | cheesy | -18.71309829 | -63.14113346 |
| Cluster-2 | caramel | -47.6863671 | 1.707411776 |
| Cluster-2 | maple | -45.01300251 | 1.051829715 |
| Cluster-2 | sugar | -46.91999995 | -1.428272404 |
| Cluster-2 | fenugreek | -43.54721027 | 0.213668944 |
| Cluster-2 | bready | -45.59643383 | -2.946699648 |
| Cluster-3 | fruity | 32.28420799 | 45.90618156 |
| Cluster-3 | sweet | 30.6136945 | 46.83344914 |
| Cluster-3 | fatty | 5.247314789 | -48.64827249 |
| Cluster-3 | floral | 30.68242658 | 50.03924369 |
| Cluster-3 | citrus | 40.82751651 | 42.17061727 |

Table S2: Detail information for t-SNE plots and odor descriptors clustering results.

| Cluster ID | Descriptor | tSNE-1 | tSNE-2 |
|---|---|---|---|
| Cluster-3 | rose | 29.54268423 | 52.92625395 |
| Cluster-3 | spicy | 26.78001678 | 39.07438665 |
| Cluster-3 | woody | 31.14014059 | 37.46076127 |
| Cluster-3 | green | 33.67742542 | 44.22180702 |
| Cluster-3 | clean | 45.33447025 | 34.57212478 |
| Cluster-3 | waxy | 6.570540198 | -47.89786453 |
| Cluster-3 | herbal | 31.16723552 | 39.77424195 |
| Cluster-3 | honey | 27.3305739 | 53.20950331 |
| Cluster-3 | fresh | 37.95444618 | 41.41137559 |
| Cluster-3 | aldehydic | 42.67318396 | 42.64007396 |
| Cluster-3 | soapy | 8.899339314 | -47.34837657 |
| Cluster-4 | nutty | 59.22022553 | -15.58122905 |
| Cluster-4 | meaty | 56.79612815 | -19.98956729 |
| Cluster-4 | sulfurous | 52.70307384 | -25.98201245 |
| Cluster-4 | coffee | 56.29849753 | -17.51490811 |
| Cluster-4 | burnt | -52.97429116 | -0.911326969 |
| Cluster-4 | roasted | 58.56101263 | -18.4645208 |
| Cluster-4 | cooked | 61.75667121 | -20.80438931 |
| Cluster-4 | meat | 63.53507415 | -20.29517537 |
| Cluster-4 | chicken | 45.78926426 | -10.42457546 |
| Cluster-4 | savory | 45.58939902 | -14.26776243 |
| Cluster-4 | lamb | 46.03675492 | -12.13056549 |
| Cluster-5 | vanilla | -10.17583701 | -51.2626144 |
| Cluster-5 | powdery | 38.34882629 | 20.64818773 |

Table S2: Detail information for t-SNE plots and odor descriptors clustering results.

| Cluster ID | Descriptor | tSNE-1 | tSNE-2 |
|---|---|---|---|
| Cluster-5 | balsam | -5.36195298 | 2.816322588 |
| Cluster-5 | cherry | -44.24624061 | -39.50670817 |
| Cluster-5 | almond | -45.79303862 | -40.12604524 |
| Cluster-5 | bitter | -47.43145886 | -40.7437799 |
| Cluster-5 | buttery | -8.812135104 | -57.69198421 |
| Cluster-5 | tobacco | -12.23321403 | -40.91425889 |
| Cluster-5 | mild | 28.12871694 | -18.85319537 |
| Cluster-5 | coumarin | -9.189469978 | -41.9533735 |
| Cluster-5 | creamy | -8.221294285 | -52.6783349 |
| Cluster-5 | coconut | -5.723264934 | -46.35066944 |
| Cluster-5 | tonka | -8.868494775 | -40.46657102 |
| Cluster-5 | dairy | -7.341449986 | -55.76946676 |
| Cluster-5 | lactonic | -4.813457384 | -44.7893436 |
| Cluster-5 | hay | -9.772780849 | -38.30496724 |
| Cluster-5 | milky | -5.806307346 | -55.46267954 |
| Cluster-5 | caramellic | -46.21964172 | 5.127307359 |
| Cluster-6 | hawthorn | -5.888671655 | -24.40186009 |
| Cluster-6 | anisic | 2.342832544 | 8.550096993 |
| Cluster-6 | anise | -1.402667592 | 9.648775902 |
| Cluster-6 | lilac | 0.109100931 | -24.19638442 |
| Cluster-6 | licorice | 0.768599124 | 12.13706781 |
| Cluster-6 | sassafrass | -0.95275496 | 11.58949243 |
| Cluster-6 | fennel | -1.794817093 | 13.58045397 |
| Cluster-6 | wintergreen | 9.20882022 | 10.50733025 |

Table S2: Detail information for t-SNE plots and odor descriptors clustering results.

| Cluster ID | Descriptor | tSNE-1 | tSNE-2 |
|---|---|---|---|
| Cluster-6 | mimosa | -7.147032128 | -24.1168678 |
| Cluster-7 | balsamic | 7.699020786 | 33.88950037 |
| Cluster-7 | mint | 11.85723369 | 12.27085378 |
| Cluster-7 | aromatic | 6.642422954 | 28.99529676 |
| Cluster-7 | cinnamyl | 4.590226761 | 31.00642554 |
| Cluster-7 | pine | 18.98823679 | 19.70705364 |
| Cluster-7 | camphor | 20.85043223 | 19.79125759 |
| Cluster-7 | incense | 15.55770909 | 27.08809268 |
| Cluster-7 | resinous | 7.110144401 | 31.61014108 |
| Cluster-7 | medical | 13.41572007 | 13.95508741 |
| Cluster-7 | terpenic | 11.41920752 | 24.74930231 |
| Cluster-7 | fir | 15.195255 | 18.50113888 |
| Cluster-7 | needle | 14.30053817 | 19.44539876 |
| Cluster-7 | labdanum | 13.56650273 | 30.84995353 |
| Cluster-7 | thujone | 11.37161181 | 19.47761628 |
| Cluster-8 | faint | 30.78751623 | -15.07500298 |
| Cluster-8 | juicy | -27.33151231 | 17.60500466 |
| Cluster-8 | winey | -50.28583828 | 35.39941721 |
| Cluster-8 | powerful | -36.61484894 | 9.42982071 |
| Cluster-8 | bland | 30.11379334 | -16.21822358 |
| Cluster-8 | cognac | -43.5492797 | 21.69485713 |
| Cluster-8 | rum | -42.13171278 | 11.42288593 |
| Cluster-8 | brown | -41.17032805 | 4.050518383 |
| Cluster-8 | ether | -39.62265632 | 8.765479313 |

Table S2: Detail information for t-SNE plots and odor descriptors clustering results.

| Cluster ID | Descriptor | tSNE-1 | tSNE-2 |
|---|---|---|---|
| Cluster-8 | estery | -25.28016981 | 15.90149478 |
| Cluster-8 | oil | -42.66969313 | 26.54273156 |
| Cluster-8 | chemical | -50.93716547 | 36.83577946 |
| Cluster-8 | and | 33.16442227 | -1.118106807 |
| Cluster-8 | alcohol | -0.638948827 | -5.250128508 |
| Cluster-8 | fusel | -44.27658852 | 25.57130111 |
| Cluster-8 | ripe | -42.63769873 | -7.328720575 |
| Cluster-8 | alcoholic | -46.09131263 | 25.86321998 |
| Cluster-8 | slightly | 33.06133083 | -3.301437571 |
| Cluster-9 | plastic | 23.52025288 | 11.43818455 |
| Cluster-9 | strong | -2.817962631 | -13.32344603 |
| Cluster-9 | sharp | 1.859400584 | -9.157832517 |
| Cluster-9 | grassy | -18.98592834 | 1.45820526 |
| Cluster-9 | narcissus | 3.352994152 | -23.97854667 |
| Cluster-9 | cortex | 4.976255314 | -24.21956565 |
| Cluster-9 | leafy | 7.010288949 | -24.78930241 |
| Cluster-9 | musty | 56.90434982 | -3.079950909 |
| Cluster-9 | earthy | 50.14166999 | -3.150031834 |
| Cluster-9 | hyacinth | 1.516351349 | -14.96990728 |
| Cluster-9 | spice | -2.302134181 | 20.14339363 |
| Cluster-9 | foliage | 0.628055196 | -13.79732734 |
| Cluster-9 | mushroom | 48.69810531 | -4.241257982 |
| Cluster-9 | geranium | 30.82666116 | 56.17875017 |
| Cluster-9 | vegetable | 53.68089287 | -30.58787858 |

Table S2: Detail information for t-SNE plots and odor descriptors clustering results.

| Cluster ID | Descriptor | tSNE-1 | tSNE-2 |
|---|---|---|---|
| Cluster-9 | red | 31.26951583 | 57.65427888 |
| Cluster-9 | metallic | 38.34751051 | -30.11163169 |
| Cluster-9 | orchid | -7.876683665 | 1.504189558 |
| Cluster-9 | weedy | -15.48615922 | 2.807765469 |
| Cluster-9 | galbanum | 10.9951667 | -6.239541243 |
| Cluster-9 | bean | -11.34889683 | 0.782636563 |
| Cluster-9 | cumin | -21.59764207 | -1.329244629 |
| Cluster-9 | mossy | 25.16980924 | 11.01725431 |
| Cluster-9 | pepper | 11.40118857 | -3.064341962 |
| Cluster-10 | berry | -46.80868885 | -11.63286657 |
| Cluster-10 | tropical | 19.63873622 | -8.265416427 |
| Cluster-10 | raspberry | -50.54324659 | -10.82966697 |
| Cluster-10 | strawberry | -49.99547071 | -7.965720578 |
| Cluster-10 | black | 13.52191316 | -13.09639395 |
| Cluster-10 | mango | 17.94135817 | -13.57156825 |
| Cluster-10 | jam | -46.51347336 | -9.119485549 |
| Cluster-10 | candy | -45.09497548 | -5.786084615 |
| Cluster-10 | passion | 15.31426343 | -10.58479476 |
| Cluster-10 | buchu | 14.83225317 | -16.27484659 |
| Cluster-10 | catty | 15.06699654 | -13.30323756 |
| Cluster-11 | grapefruit | -20.96784154 | 41.96260602 |
| Cluster-11 | orange | -29.24988443 | 25.04055319 |
| Cluster-11 | petitgrain | -24.50941611 | 29.73374341 |
| Cluster-11 | lemon | 16.5508425 | 2.746692048 |

Table S2: Detail information for t-SNE plots and odor descriptors clustering results.

| Cluster ID | Descriptor | tSNE-1 | tSNE-2 |
|---|---|---|---|
| Cluster-11 | peel | -22.75728034 | 40.52555781 |
| Cluster-11 | lime | 18.35203124 | 2.500003431 |
| Cluster-11 | mandarin | -27.98436743 | 36.12723491 |
| Cluster-11 | blossom | -30.52972302 | 25.9975111 |
| Cluster-11 | neroli | -25.99689902 | 29.39080792 |
| Cluster-11 | cologne | 20.191318 | 2.736239593 |
| Cluster-11 | tangerine | -27.84443987 | 37.51406965 |
| Cluster-12 | walnut | -35.99947812 | -37.55052501 |
| Cluster-12 | fermented | -48.15797706 | 26.82605574 |
| Cluster-12 | skin | -35.24897841 | -29.42860771 |
| Cluster-12 | chocolate | 59.73813495 | -4.65827757 |
| Cluster-12 | cocoa | 61.01028689 | -5.553424625 |
| Cluster-12 | peanut | -30.13398858 | -37.57407778 |
| Cluster-12 | potato | -26.5043347 | -37.07277082 |
| Cluster-12 | nut | -34.24507757 | -30.46461554 |
| Cluster-12 | popcorn | -26.34537558 | -30.37641429 |
| Cluster-12 | beef | 49.17296054 | -11.93038363 |
| Cluster-12 | corn | -27.8437505 | -33.46855408 |
| Cluster-12 | chip | -26.48754411 | -32.62068762 |
| Cluster-12 | baked | -24.51503303 | -39.87733997 |
| Cluster-12 | hazelnut | -31.3209417 | -35.37547279 |
| Cluster-12 | toasted | -21.4142279 | -32.35514864 |
| Cluster-12 | grain | -22.8950864 | -32.05722548 |
| Cluster-13 | phenolic | -62.42485117 | 38.75057865 |

Table S2: Detail information for t-SNE plots and odor descriptors clustering results.

| Cluster ID | Descriptor | tSNE-1 | tSNE-2 |
|---|---|---|---|
| Cluster-13 | solvent | -27.45323318 | -4.288855126 |
| Cluster-13 | herbaceous | -17.22070744 | -14.31484944 |
| Cluster-13 | minty | -21.41686624 | -11.44930605 |
| Cluster-13 | camphoreous | -9.70013315 | -8.901471238 |
| Cluster-13 | rosemary | -11.94202908 | -7.203191643 |
| Cluster-13 | medicinal | -61.02370514 | 38.04184184 |
| Cluster-13 | camphoraceous | -27.72207632 | -13.1349499 |
| Cluster-13 | smoky | -55.00811064 | -0.176999702 |
| Cluster-13 | cool | -23.4943 | -16.66210051 |
| Cluster-13 | celery | -32.41200316 | -2.221053032 |
| Cluster-13 | caraway | -24.38853675 | -5.019965202 |
| Cluster-13 | mentholic | -23.52421837 | -13.645552 |
| Cluster-13 | spearmint | -24.57391752 | -6.896377548 |
| Cluster-13 | peppery | 1.928784116 | 1.047260901 |
| Cluster-13 | terpene | 3.381102925 | 1.468807937 |
| Cluster-13 | eucalyptus | -10.01437278 | -7.241365116 |
| Cluster-13 | peppermint | -24.87916854 | -14.35079675 |
| Cluster-13 | cooling | -24.35726501 | -11.16783113 |
| Cluster-14 | alliaceous | 47.45388915 | -26.40825159 |
| Cluster-14 | onion | 50.33200201 | -26.09990545 |
| Cluster-14 | garlic | 49.19662044 | -27.68063127 |
| Cluster-14 | horseradish | 49.57880532 | -35.49218908 |
| Cluster-14 | sulfury | 50.3767848 | -23.37192614 |
| Cluster-14 | tomato | 41.76063671 | -31.33803447 |

Table S2: Detail information for t-SNE plots and odor descriptors clustering results.

| Cluster ID | Descriptor | tSNE-1 | tSNE-2 |
|---|---|---|---|
| Cluster-14 | chrysanthemum | 43.46929486 | -35.47641761 |
| Cluster-14 | pungent | 50.57930547 | -39.13163512 |
| Cluster-14 | cabbage | 46.43746864 | -31.91796257 |
| Cluster-14 | mustard | 49.60661666 | -36.91778314 |
| Cluster-14 | radish | 44.45057298 | -33.47380264 |
| Cluster-15 | ethereal | -53.31311038 | 15.22278107 |
| Cluster-15 | wine | -45.56334001 | 15.10311041 |
| Cluster-15 | apricot | -65.15574543 | -9.942602568 |
| Cluster-15 | plum | -63.33586731 | -7.73439385 |
| Cluster-15 | banana | -56.8621136 | 14.69876279 |
| Cluster-15 | peach | -65.62687488 | -11.18080326 |
| Cluster-15 | pineapple | -59.13641095 | 16.02716176 |
| Cluster-15 | apple | -57.89915901 | 17.68630994 |
| Cluster-15 | pear | -58.47414685 | 19.40970368 |
| Cluster-15 | wine-like | -53.67146865 | 18.0767722 |
| Cluster-15 | grape | -33.85279268 | 25.32810299 |
| Cluster-15 | brandy | -44.26774236 | 13.58621323 |
| Cluster-16 | oily | 2.156074925 | -47.60054747 |
| Cluster-16 | rhubarb | -7.111432064 | 53.45390837 |
| Cluster-16 | natural | 3.167692298 | 48.00587192 |
| Cluster-16 | jasmin | -7.277409901 | 43.51688084 |
| Cluster-16 | lily | 27.05693597 | -39.58942662 |
| Cluster-16 | jasmine | -42.31589445 | -16.71057486 |
| Cluster-16 | valley | -22.27081873 | 14.36067767 |

Table S2: Detail information for t-SNE plots and odor descriptors clustering results.

| Cluster ID | Descriptor | tSNE-1 | tSNE-2 |
|---|---|---|---|
| Cluster-16 | ylang | -5.036142738 | 46.81993013 |
| Cluster-16 | petal | -1.89428447 | 48.9010504 |
| Cluster-16 | ambrette | 34.03072578 | 19.86809085 |
| Cluster-16 | flower | -20.98442744 | 13.54488991 |
| Cluster-16 | gardenia | -6.470521383 | 51.02097919 |
| Cluster-16 | magnolia | 27.3346825 | -37.21044915 |
| Cluster-16 | muguet | 25.3018962 | -39.66717327 |
| Cluster-16 | tuberose | -6.07879514 | 47.93456174 |
| Cluster-16 | orange flower | -8.811580304 | 47.71642281 |
| Cluster-16 | seed | 31.71695375 | 21.1440193 |
| Cluster-17 | violet | 17.64468675 | -30.64132027 |
| Cluster-17 | cucumber | 14.44404581 | -36.85908054 |
| Cluster-17 | orris | 17.02059101 | -29.20502834 |
| Cluster-17 | leaf | 19.21165398 | -32.37526793 |
| Cluster-17 | melon | 13.36196027 | -37.57718839 |
| Cluster-17 | watery | 49.55693176 | 34.26375536 |
| Cluster-17 | rind | -29.17153621 | 40.21500345 |
| Cluster-17 | ozone | 49.48473473 | 32.04521718 |
| Cluster-17 | marine | 49.94978551 | 30.57703188 |
| Cluster-18 | dry | 34.28063971 | 14.74128223 |
| Cluster-18 | musk | 36.69153738 | 18.37994002 |
| Cluster-18 | civet | 41.47794693 | 14.08441206 |
| Cluster-18 | animal | 42.41504427 | 16.25245273 |
| Cluster-18 | cedar | 32.17724013 | 9.238710459 |

Table S2: Detail information for t-SNE plots and odor descriptors clustering results.

| Cluster ID | Descriptor | tSNE-1 | tSNE-2 |
| --- | --- | --- | --- |
| Cluster-18 | sandalwood | 25.76937053 | 4.507027384 |
| Cluster-18 | amber | 36.08647378 | 16.44850253 |
| Cluster-18 | greasy | 50.59977229 | 27.82859406 |
| Cluster-18 | vetiver | 32.94906582 | 4.64196648 |
| Cluster-18 | dusty | 36.66868104 | 11.09196856 |
| Cluster-18 | ambergris | 33.82872679 | 10.42531764 |
| Cluster-18 | leather | 44.49831521 | 17.0605652 |
| Cluster-18 | wood | 17.21674257 | 31.01732186 |
| Cluster-18 | patchouli | 29.65299656 | 8.705027041 |
| Cluster-18 | acetate | 32.89728136 | 2.891643927 |
| Cluster-18 | old | 16.01542784 | 30.2451427 |
| Cluster-19 | warm | 23.85259765 | 36.81536529 |
| Cluster-19 | cinnamon | -0.367667318 | 29.14826097 |
| Cluster-19 | cassia | -0.215046385 | 30.58575749 |
| Cluster-19 | deep | -2.092695659 | 32.55760265 |
| Cluster-19 | clove | 22.19422231 | 39.81564284 |
| Cluster-19 | carnation | 21.04503392 | 39.05199104 |
| Cluster-19 | root | -6.560638159 | 12.23436958 |
| Cluster-19 | bay | -11.98181982 | 12.74258117 |
| Cluster-19 | ginger | -10.53855636 | 12.33340318 |
| Cluster-20 | tea | 4.50103649 | 20.96389898 |
| Cluster-20 | parsley | -15.06539715 | -5.947100256 |
| Cluster-20 | chamomile | 3.10789135 | 18.6832698 |
| Cluster-20 | lavender | -19.12237445 | 30.73651675 |

Table S2: Detail information for t-SNE plots and odor descriptors clustering results.

| Cluster ID | Descriptor | tSNE-1 | tSNE-2 |
|---|---|---|---|
| Cluster-20 | rooty | -8.069397007 | 23.20435185 |
| Cluster-20 | bergamot | -18.89362739 | 28.97640077 |
| Cluster-20 | sage | -17.34670015 | 25.6707589 |
| Cluster-20 | basil | 1.383777582 | 17.07038738 |
| Cluster-20 | bois | -10.57844031 | 22.01014207 |
| Cluster-20 | clary | -16.05327711 | 25.59255282 |
| Cluster-20 | blueberry | -11.62759784 | 21.03669983 |
| Cluster-20 | thyme | -18.73775464 | -4.123075919 |

Table S3: Correlations of Word2Vec embedded vectors for odor categories.

| Cluster-ID | Correlation | Standard deviation | Cluster-ID | Correlation | Standard deviation |
|---|---|---|---|---|---|
| Cluster-1 | 0.258 | 0.119 | Cluster-11 | 0.323 | 0.130 |
| Cluster-2 | 0.382 | 0.099 | Cluster-12 | 0.265 | 0.130 |
| Cluster-3 | 0.247 | 0.123 | Cluster-13 | 0.324 | 0.165 |
| Cluster-4 | 0.310 | 0.152 | Cluster-14 | 0.479 | 0.128 |
| Cluster-5 | 0.288 | 0.150 | Cluster-15 | 0.401 | 0.176 |
| Cluster-6 | 0.443 | 0.067 | Cluster-16 | 0.320 | 0.171 |
| Cluster-7 | 0.245 | 0.168 | Cluster-17 | 0.270 | 0.158 |
| Cluster-8 | 0.213 | 0.165 | Cluster-18 | 0.231 | 0.129 |
| Cluster-9 | 0.217 | 0.139 | Cluster-19 | 0.227 | 0.174 |
| Cluster-10 | 0.187 | 0.136 | Cluster-20 | 0.240 | 0.083 |

Table S4: Odor cluster identification results by GLVQ and GBDT models.

(a) Alexnet results

| Model | GLVQ | | | | GDBT | | | |
|---|---|---|---|---|---|---|---|---|
| Cluster ID | ROC | Precision | Recall | F-score | ROC | Precision | Recall | F-score |
| Cluster-1 | 0.668 | 0.526 | 0.611 | 0.433 | 0.720 | 0.536 | 0.639 | 0.482 |
| Cluster-2 | 0.824 | 0.522 | 0.699 | 0.423 | 0.849 | 0.538 | 0.786 | 0.493 |
| Cluster-3 | 0.711 | 0.651 | 0.660 | 0.655 | 0.794 | 0.680 | 0.672 | 0.676 |
| Cluster-4 | 0.790 | 0.670 | 0.748 | 0.689 | 0.873 | 0.715 | 0.794 | 0.740 |
| Cluster-5 | 0.652 | 0.570 | 0.609 | 0.563 | 0.697 | 0.586 | 0.648 | 0.557 |
| Cluster-6 | 0.779 | 0.529 | 0.690 | 0.438 | 0.856 | 0.557 | 0.780 | 0.541 |
| Cluster-7 | 0.712 | 0.540 | 0.661 | 0.479 | 0.758 | 0.552 | 0.689 | 0.522 |
| Cluster-8 | 0.693 | 0.526 | 0.571 | 0.522 | 0.758 | 0.542 | 0.695 | 0.502 |
| Cluster-9 | 0.589 | 0.543 | 0.557 | 0.526 | 0.659 | 0.586 | 0.615 | 0.565 |
| Cluster-10 | 0.640 | 0.555 | 0.603 | 0.506 | 0.675 | 0.579 | 0.647 | 0.542 |
| Cluster-11 | 0.661 | 0.536 | 0.629 | 0.476 | 0.773 | 0.555 | 0.687 | 0.512 |
| Cluster-12 | 0.697 | 0.545 | 0.650 | 0.508 | 0.718 | 0.547 | 0.651 | 0.517 |
| Cluster-13 | 0.745 | 0.575 | 0.670 | 0.448 | 0.814 | 0.597 | 0.718 | 0.576 |
| Cluster-14 | 0.784 | 0.584 | 0.730 | 0.580 | 0.878 | 0.659 | 0.801 | 0.695 |
| Cluster-15 | 0.745 | 0.608 | 0.666 | 0.559 | 0.820 | 0.662 | 0.745 | 0.657 |
| Cluster-16 | 0.615 | 0.558 | 0.611 | 0.526 | 0.722 | 0.574 | 0.646 | 0.534 |
| Cluster-17 | 0.599 | 0.516 | 0.565 | 0.429 | 0.707 | 0.536 | 0.629 | 0.491 |
| Cluster-18 | 0.762 | 0.596 | 0.718 | 0.599 | 0.838 | 0.604 | 0.742 | 0.605 |
| Cluster-19 | 0.677 | 0.520 | 0.611 | 0.402 | 0.794 | 0.538 | 0.695 | 0.480 |
| Cluster-20 | 0.734 | 0.525 | 0.637 | 0.387 | 0.762 | 0.534 | 0.681 | 0.462 |

Table S4: Odor cluster identification results by GLVQ and GBDT models.

(b) Densenet results

| Model | GLVQ | | | | GDBT | | | |
|---|---|---|---|---|---|---|---|---|
| Cluster ID | ROC | Precision | Recall | F-score | ROC | Precision | Recall | F-score |
| Cluster-1 | 0.667 | 0.529 | 0.621 | 0.434 | 0.756 | 0.546 | 0.675 | 0.505 |
| Cluster-2 | 0.865 | 0.531 | 0.766 | 0.448 | 0.895 | 0.548 | 0.815 | 0.527 |
| Cluster-3 | 0.683 | 0.608 | 0.660 | 0.607 | 0.818 | 0.709 | 0.691 | 0.699 |
| Cluster-4 | 0.813 | 0.685 | 0.750 | 0.705 | 0.878 | 0.732 | 0.816 | 0.759 |
| Cluster-5 | 0.655 | 0.562 | 0.606 | 0.531 | 0.707 | 0.594 | 0.656 | 0.580 |
| Cluster-6 | 0.809 | 0.528 | 0.682 | 0.380 | 0.888 | 0.558 | 0.815 | 0.531 |
| Cluster-7 | 0.707 | 0.541 | 0.670 | 0.451 | 0.766 | 0.558 | 0.721 | 0.520 |
| Cluster-8 | 0.727 | 0.549 | 0.650 | 0.552 | 0.804 | 0.551 | 0.743 | 0.506 |
| Cluster-9 | 0.620 | 0.562 | 0.582 | 0.511 | 0.688 | 0.598 | 0.632 | 0.567 |
| Cluster-10 | 0.626 | 0.556 | 0.602 | 0.533 | 0.692 | 0.573 | 0.636 | 0.541 |
| Cluster-11 | 0.740 | 0.553 | 0.683 | 0.506 | 0.799 | 0.565 | 0.709 | 0.539 |
| Cluster-12 | 0.694 | 0.547 | 0.651 | 0.517 | 0.741 | 0.553 | 0.667 | 0.529 |
| Cluster-13 | 0.743 | 0.582 | 0.695 | 0.503 | 0.839 | 0.607 | 0.740 | 0.590 |
| Cluster-14 | 0.784 | 0.633 | 0.772 | 0.661 | 0.874 | 0.638 | 0.789 | 0.669 |
| Cluster-15 | 0.735 | 0.621 | 0.684 | 0.554 | 0.844 | 0.672 | 0.754 | 0.674 |
| Cluster-16 | 0.628 | 0.545 | 0.586 | 0.518 | 0.723 | 0.590 | 0.677 | 0.559 |
| Cluster-17 | 0.655 | 0.531 | 0.620 | 0.464 | 0.746 | 0.557 | 0.693 | 0.533 |
| Cluster-18 | 0.773 | 0.592 | 0.716 | 0.589 | 0.847 | 0.609 | 0.740 | 0.618 |
| Cluster-19 | 0.714 | 0.528 | 0.658 | 0.397 | 0.794 | 0.543 | 0.726 | 0.487 |
| Cluster-20 | 0.710 | 0.523 | 0.618 | 0.336 | 0.811 | 0.551 | 0.745 | 0.509 |

Table S4: Odor cluster identification results by GLVQ and GBDT models.

(c) Resnet results

| Model | GLVQ | | | | GDBT | | | |
|---|---|---|---|---|---|---|---|---|
| Cluster ID | ROC | Precision | Recall | F-score | ROC | Precision | Recall | F-score |
| Cluster-1 | 0.693 | 0.531 | 0.627 | 0.451 | 0.753 | 0.547 | 0.678 | 0.508 |
| Cluster-2 | 0.838 | 0.531 | 0.765 | 0.447 | 0.883 | 0.540 | 0.810 | 0.496 |
| Cluster-3 | 0.706 | 0.636 | 0.676 | 0.646 | 0.786 | 0.694 | 0.662 | 0.675 |
| Cluster-4 | 0.820 | 0.693 | 0.775 | 0.716 | 0.864 | 0.698 | 0.792 | 0.723 |
| Cluster-5 | 0.650 | 0.572 | 0.625 | 0.531 | 0.700 | 0.578 | 0.633 | 0.551 |
| Cluster-6 | 0.762 | 0.524 | 0.656 | 0.401 | 0.854 | 0.549 | 0.773 | 0.514 |
| Cluster-7 | 0.713 | 0.539 | 0.654 | 0.481 | 0.766 | 0.548 | 0.681 | 0.512 |
| Cluster-8 | 0.622 | 0.526 | 0.571 | 0.523 | 0.739 | 0.531 | 0.640 | 0.489 |
| Cluster-9 | 0.624 | 0.572 | 0.595 | 0.514 | 0.660 | 0.566 | 0.589 | 0.536 |
| Cluster-10 | 0.642 | 0.557 | 0.606 | 0.514 | 0.686 | 0.577 | 0.642 | 0.545 |
| Cluster-11 | 0.716 | 0.550 | 0.678 | 0.484 | 0.779 | 0.556 | 0.693 | 0.510 |
| Cluster-12 | 0.712 | 0.550 | 0.649 | 0.532 | 0.725 | 0.549 | 0.662 | 0.515 |
| Cluster-13 | 0.746 | 0.583 | 0.699 | 0.509 | 0.815 | 0.595 | 0.711 | 0.577 |
| Cluster-14 | 0.825 | 0.627 | 0.787 | 0.652 | 0.865 | 0.617 | 0.783 | 0.635 |
| Cluster-15 | 0.764 | 0.632 | 0.705 | 0.597 | 0.848 | 0.680 | 0.767 | 0.682 |
| Cluster-16 | 0.658 | 0.567 | 0.625 | 0.547 | 0.721 | 0.593 | 0.681 | 0.561 |
| Cluster-17 | 0.665 | 0.527 | 0.608 | 0.433 | 0.734 | 0.551 | 0.680 | 0.516 |
| Cluster-18 | 0.748 | 0.619 | 0.734 | 0.638 | 0.818 | 0.600 | 0.733 | 0.600 |
| Cluster-19 | 0.697 | 0.532 | 0.679 | 0.421 | 0.790 | 0.543 | 0.719 | 0.493 |
| Cluster-20 | 0.670 | 0.521 | 0.610 | 0.436 | 0.718 | 0.522 | 0.608 | 0.469 |

Table S4: Odor cluster identification results by GLVQ and GBDT models.

(d) VGG results

| Model | GLVQ | | | | GDBT | | | |
|---|---|---|---|---|---|---|---|---|
| Cluster ID | ROC | Precision | Recall | F-score | ROC | Precision | Recall | F-score |
| Cluster-1 | 0.642 | 0.517 | 0.572 | 0.368 | 0.714 | 0.540 | 0.652 | 0.498 |
| Cluster-2 | 0.858 | 0.526 | 0.732 | 0.413 | 0.864 | 0.541 | 0.813 | 0.499 |
| Cluster-3 | 0.679 | 0.599 | 0.643 | 0.598 | 0.803 | 0.705 | 0.674 | 0.687 |
| Cluster-4 | 0.757 | 0.600 | 0.680 | 0.581 | 0.866 | 0.677 | 0.776 | 0.696 |
| Cluster-5 | 0.639 | 0.557 | 0.595 | 0.535 | 0.715 | 0.594 | 0.654 | 0.583 |
| Cluster-6 | 0.786 | 0.537 | 0.739 | 0.442 | 0.883 | 0.557 | 0.813 | 0.528 |
| Cluster-7 | 0.724 | 0.539 | 0.666 | 0.425 | 0.766 | 0.556 | 0.717 | 0.516 |
| Cluster-8 | 0.579 | 0.527 | 0.579 | 0.521 | 0.805 | 0.551 | 0.743 | 0.506 |
| Cluster-9 | 0.593 | 0.541 | 0.553 | 0.478 | 0.649 | 0.568 | 0.592 | 0.544 |
| Cluster-10 | 0.655 | 0.555 | 0.604 | 0.488 | 0.700 | 0.585 | 0.659 | 0.546 |
| Cluster-11 | 0.706 | 0.545 | 0.659 | 0.487 | 0.767 | 0.550 | 0.664 | 0.517 |
| Cluster-12 | 0.646 | 0.532 | 0.617 | 0.454 | 0.700 | 0.545 | 0.649 | 0.507 |
| Cluster-13 | 0.723 | 0.585 | 0.695 | 0.556 | 0.831 | 0.608 | 0.741 | 0.592 |
| Cluster-14 | 0.763 | 0.561 | 0.693 | 0.515 | 0.871 | 0.612 | 0.767 | 0.629 |
| Cluster-15 | 0.693 | 0.608 | 0.665 | 0.543 | 0.821 | 0.679 | 0.773 | 0.673 |
| Cluster-16 | 0.581 | 0.552 | 0.576 | 0.554 | 0.707 | 0.578 | 0.654 | 0.537 |
| Cluster-17 | 0.646 | 0.528 | 0.612 | 0.446 | 0.750 | 0.553 | 0.688 | 0.517 |
| Cluster-18 | 0.770 | 0.593 | 0.728 | 0.585 | 0.815 | 0.610 | 0.757 | 0.614 |
| Cluster-19 | 0.714 | 0.521 | 0.615 | 0.364 | 0.824 | 0.547 | 0.749 | 0.485 |
| Cluster-20 | 0.660 | 0.512 | 0.566 | 0.415 | 0.712 | 0.534 | 0.664 | 0.482 |

Table S4: Odor cluster identification results by GLVQ and GBDT models.

(e) Molecular parameter results

| Model | GLVQ | | | | GDBT | | | |
|---|---|---|---|---|---|---|---|---|
| Cluster ID | ROC | Precision | Recall | F-score | ROC | Precision | Recall | F-score |
| Cluster-1 | 0.520 | 0.506 | 0.527 | 0.358 | 0.567 | 0.508 | 0.534 | 0.386 |
| Cluster-2 | 0.429 | 0.496 | 0.465 | 0.344 | 0.428 | 0.491 | 0.417 | 0.341 |
| Cluster-3 | 0.473 | 0.480 | 0.468 | 0.439 | 0.536 | 0.534 | 0.519 | 0.515 |
| Cluster-4 | 0.474 | 0.498 | 0.497 | 0.398 | 0.512 | 0.504 | 0.506 | 0.427 |
| Cluster-5 | 0.519 | 0.513 | 0.523 | 0.437 | 0.585 | 0.542 | 0.572 | 0.477 |
| Cluster-6 | 0.523 | 0.499 | 0.492 | 0.386 | 0.587 | 0.514 | 0.595 | 0.401 |
| Cluster-7 | 0.531 | 0.500 | 0.498 | 0.375 | 0.625 | 0.518 | 0.576 | 0.413 |
| Cluster-8 | 0.483 | 0.491 | 0.453 | 0.377 | 0.560 | 0.506 | 0.531 | 0.374 |
| Cluster-9 | 0.420 | 0.458 | 0.444 | 0.424 | 0.532 | 0.500 | 0.499 | 0.437 |
| Cluster-10 | 0.521 | 0.522 | 0.541 | 0.454 | 0.554 | 0.521 | 0.539 | 0.438 |
| Cluster-11 | 0.477 | 0.497 | 0.487 | 0.366 | 0.470 | 0.492 | 0.470 | 0.371 |
| Cluster-12 | 0.492 | 0.499 | 0.496 | 0.406 | 0.590 | 0.522 | 0.582 | 0.431 |
| Cluster-13 | 0.488 | 0.492 | 0.481 | 0.392 | 0.513 | 0.500 | 0.499 | 0.388 |
| Cluster-14 | 0.567 | 0.511 | 0.537 | 0.407 | 0.623 | 0.536 | 0.619 | 0.442 |
| Cluster-15 | 0.538 | 0.519 | 0.527 | 0.504 | 0.541 | 0.520 | 0.531 | 0.467 |
| Cluster-16 | 0.474 | 0.473 | 0.459 | 0.458 | 0.483 | 0.503 | 0.506 | 0.428 |
| Cluster-17 | 0.483 | 0.498 | 0.492 | 0.437 | 0.574 | 0.517 | 0.566 | 0.386 |
| Cluster-18 | 0.491 | 0.499 | 0.498 | 0.394 | 0.530 | 0.515 | 0.540 | 0.408 |
| Cluster-19 | 0.522 | 0.506 | 0.534 | 0.410 | 0.528 | 0.500 | 0.500 | 0.350 |
| Cluster-20 | 0.416 | 0.492 | 0.458 | 0.398 | 0.431 | 0.498 | 0.488 | 0.371 |

Table S4: Odor cluster identification results by GLVQ and GBDT models.

(f) Finger print results

| Model | GLVQ | | | | GDBT | | | |
|---|---|---|---|---|---|---|---|---|
| Cluster ID | ROC | Precision | Recall | F-score | ROC | Precision | Recall | F-score |
| Cluster-1 | 0.591 | 0.514 | 0.556 | 0.436 | 0.590 | 0.518 | 0.571 | 0.444 |
| Cluster-2 | 0.395 | 0.494 | 0.455 | 0.228 | 0.429 | 0.492 | 0.429 | 0.331 |
| Cluster-3 | 0.528 | 0.517 | 0.526 | 0.500 | 0.545 | 0.535 | 0.539 | 0.536 |
| Cluster-4 | 0.485 | 0.498 | 0.497 | 0.376 | 0.544 | 0.525 | 0.547 | 0.464 |
| Cluster-5 | 0.529 | 0.519 | 0.525 | 0.516 | 0.572 | 0.528 | 0.549 | 0.471 |
| Cluster-6 | 0.535 | 0.500 | 0.499 | 0.443 | 0.548 | 0.504 | 0.528 | 0.379 |
| Cluster-7 | 0.548 | 0.508 | 0.527 | 0.290 | 0.555 | 0.517 | 0.571 | 0.387 |
| Cluster-8 | 0.484 | 0.503 | 0.514 | 0.444 | 0.528 | 0.506 | 0.531 | 0.398 |
| Cluster-9 | 0.513 | 0.513 | 0.515 | 0.416 | 0.537 | 0.520 | 0.526 | 0.477 |
| Cluster-10 | 0.499 | 0.499 | 0.498 | 0.415 | 0.596 | 0.532 | 0.560 | 0.454 |
| Cluster-11 | 0.516 | 0.503 | 0.509 | 0.434 | 0.536 | 0.502 | 0.508 | 0.381 |
| Cluster-12 | 0.577 | 0.515 | 0.555 | 0.375 | 0.590 | 0.515 | 0.556 | 0.426 |
| Cluster-13 | 0.485 | 0.495 | 0.489 | 0.468 | 0.480 | 0.499 | 0.497 | 0.414 |
| Cluster-14 | 0.498 | 0.501 | 0.502 | 0.288 | 0.537 | 0.506 | 0.520 | 0.402 |
| Cluster-15 | 0.528 | 0.504 | 0.507 | 0.469 | 0.550 | 0.522 | 0.534 | 0.451 |
| Cluster-16 | 0.505 | 0.506 | 0.513 | 0.436 | 0.483 | 0.495 | 0.490 | 0.432 |
| Cluster-17 | 0.489 | 0.501 | 0.502 | 0.471 | 0.495 | 0.493 | 0.471 | 0.403 |
| Cluster-18 | 0.523 | 0.504 | 0.508 | 0.485 | 0.574 | 0.517 | 0.546 | 0.442 |
| Cluster-19 | 0.550 | 0.513 | 0.548 | 0.227 | 0.576 | 0.510 | 0.558 | 0.358 |
| Cluster-20 | 0.648 | 0.516 | 0.581 | 0.441 | 0.608 | 0.514 | 0.575 | 0.373 |
